# Cis-regulatory analysis of Onecut1 expression in fate-restricted retinal progenitor cells

**DOI:** 10.1101/854778

**Authors:** Sruti Patoori, Nathalie Jean-Charles, Ariana Gopal, Sacha Sulaiman, Sneha Gopal, Brian Wang, Benjamin Souferi, Mark M. Emerson

## Abstract

**Background:** The vertebrate retina consists of six major classes of neuronal cells. During development, these cells are generated from a pool of multipotent retinal progenitor cells (RPCs) that express the gene Vsx2. Fate-restricted RPCs have recently been identified, with limited mitotic potential and cell fate possibilities compared to multipotent RPCs. One population of fate-restricted RPCs, marked by activity of the regulatory element ThrbCRM1, gives rise to both cone photoreceptors and horizontal cells. These cells do not express Vsx2, but co-express the transcription factors (TFs) Onecut1 and Otx2, which bind to ThrbCRM1. The components of the gene regulatory networks that control the transition from multipotent to fate-restricted gene expression are not known. This work aims to identify and evaluate cis-regulatory elements proximal to Onecut1 to identify the gene regulatory networks involved in RPC fate-restriction.

**Method:** We identified regulatory elements through ATAC-seq and conservation, followed by reporter assays to screen for activity based on temporal and spatial criteria. The regulatory elements of interest were subject to deletion and mutation analysis to identify functional sequences and evaluated by quantitative flow cytometry assays. Finally, we combined the enhancer::reporter assays with candidate TF overexpression to evaluate the relationship between the TFs, the enhancers, and early vertebrate retinal development. Statistical tests included ANOVA, Kruskal-Wallis, or unpaired t-tests.

**Results:** Two regulatory elements, ECR9 and ECR65, were identified to be active in ThrbCRM1(+) restricted RPCs. Candidate bHLH binding sites were identified as critical sequences in both elements. Overexpression of candidate bHLH TFs revealed specific enhancer-bHLH interactions. Nhlh1 overexpression expanded ECR65 activity into the Vsx2(+) RPC population, and overexpression of NeuroD1/NeuroG2/NeuroD4 had a similar effect on ECR9. Furthermore, bHLHs that were able to activate ectopic ECR9 reporter were able to induce endogenous Otx2 expression.

**Conclusions:** This work reports a large-scale screen to identify spatiotemporally specific regulatory elements near the Onecut1 locus. These elements were used to identify distinct populations in the developing retina. In addition, fate-restricted regulatory elements responded differentially to bHLH factors, and suggest a role for retinal bHLHs upstream of the Otx2 and Onecut1 genes during the formation of restricted RPCs from multipotent RPCs.

## Introduction

The vertebrate retina is comprised of six main classes of neuronal cells and one class of glial cells, organized into three discrete nuclear layers and two plexiform layers. These morphologically and functionally diverse cells have been characterized in multiple vertebrate species to originate from multipotent retinal progenitor cells (RPCs) (Turner and Cepko, 1987; Fekete et al., 1994). The vertebrate retina is therefore a valuable model to study neuronal cell fate choice. The process of retinal development is highly conserved throughout the vertebrate subphylum, with regards to the birth order of the various cell types (Young, 1985; Wong and Rapaport, 2009) and the developmental regulatory networks involved. However, the gene regulatory networks (GRNs) that mediate the generation of specific restricted RPCs from multipotent RPCs are largely unknown, as are the networks that function in restricted RPCs to define their fate potential.

One restricted RPC type that has been identified across zebrafish, chick, and mouse models preferentially generates cones and horizontal cells (HCs) and has been identified in zebrafish and chick through regulatory elements associated with the Thrb and Olig2 genes (Emerson et al., 2013; Hafler et al., 2012; Suzuki et al., 2013). For example, analysis in zebrafish and mouse RPCs showed that the same RPC can give rise to both cone and horizontal cell precursor cells (Hafler et al., 2012; Suzuki et al., 2013). Endogenous Thrb expression has been observed in Otx2-expressing early RPCs in the chick, suggesting that reporters driven by regulatory elements correspond to *in vivo* regulatory events in the retina (Trimarchi et al., 2008). While Otx2 expression is involved in multiple cell fates during retinal development, it has been shown that the combination of Otx2 and Onecut1 activates ThrbCRM1, which is a specific Thrb cis-regulatory element (CRE) active in cone/HC restricted RPCs (RPC[CH]) (Emerson et al., 2013). Loss-of-function mutations in Otx2 and Onecut1 affect early cone gene expression, cone number, cone type, and horizontal cell genesis (Nishida et al., 2003; Sapkota et al., 2014; Wu et al., 2013), suggesting that these transcription factors (TFs) are critical in the gene regulatory networks of ThrbCRM1 restricted RPCs.

The population of restricted RPCs marked by ThrbCRM1 have been shown to be molecularly distinct from multipotent RPCs. ThrbCRM1(+) RPCs downregulate multipotent RPC genes such as Vsx2 while Onecut1 is upregulated (Buenaventura et al., 2018). Onecut1 expression is further upregulated in the HC progeny of these cells but is downregulated in the cone photoreceptor progeny. However, it is not known how Onecut1 expression is activated in the ThrbCRM1 RPC population or what distinguishes it from other Onecut1(+) cell populations.

The regulatory module that connects Otx2 and Onecut1 to Thrb expression demonstrates the importance of both cis- and trans-regulatory elements in directing retinal cell fate. Cell fate specification and fate restriction require the combinatorial expression of multiple developmental transcription factors. As such, cell-type specific cis-regulatory elements can define the intermediate/restricted RPCs that may be difficult to identify through only the transient expression of developmental transcription factors that are often involved in the specification of multiple retinal cell types. These regulatory elements can be used to facilitate imaging, lineage tracing, and molecular analysis, while also providing insights into the relationships between RPC populations.

We sought to identify the cis-regulatory elements upstream of Onecut1 expression in cone/HC restricted RPCs and to determine the transcription factors that occupy these elements. To this end, we conducted a multi-step screen to identify Onecut1-associated non-coding DNA elements capable of driving reporter transcription in early retinal RPCs that give rise to cones and horizontal cells in the early embryonic chick retina. The candidate regulatory elements that emerged from the screen were then bioinformatically analyzed for transcription factor binding sites. Mutational analyses facilitated the functional evaluation of these predicted TF binding sites and overexpression experiments were used to determine the relationship between the predicted transcription factors and Onecut1 expression. We identified two regulatory elements, ECR9 and ECR65, upstream of the Onecut1 coding region. These both contain predicted binding sites for bHLH transcription factors, which are known to be functionally important for retinal development. We show that both of these elements require the predicted bHLH binding sites for their activity and that each element responds to distinct bHLH factors, the transcripts of which are enriched or present in the ThrbCRM1 population. Finally, we show that NeuroD1, NeuroG2, and NeuroD4 are sufficient to induce expression of Otx2 and that all four TFs including Nhlh1 are able to induce the activity of their corresponding regulatory elements in Vsx2(+) multipotent RPCs. Ultimately, this work further clarifies components of the gene regulatory network leading to the early retinal cell fates of cone photoreceptors, horizontal cells, and retinal ganglion cells.

## Results

### Regulatory Element Identification

Two methods were employed to identify candidate cis-regulatory elements for Onecut1 (Figure 1A,B). We first examined the intergenic region 5’ of the chicken Onecut1 coding region and 3’ of WDR72, as well as a short stretch of the intergenic region 3’ of the Onecut1 coding region for evolutionarily conserved regions (ECRs). As not all cis-regulatory sequences are strongly conserved and as ECRBrowser (Ovcharenko et al., 2004) utilizes an older chick genome assembly, we also used chromatin accessibility as a means of candidate enhancer identification. Chick E5 retinae were electroporated with ThrbCRM1::GFP and UbiqC::TdT, cultured for 18-22 hours *ex vivo* and sorted into two populations: ThrbCRM1(+) cells, which were marked by both reporters and ThrbCRM1(-), which were marked by only TdT. These cells were then processed for ATAC-seq (Buenrostro et al., 2013), aligned against the galGal5 assembly and the data was visualized in the UCSC Genome Browser (Kent et al., 2002) to identify accessible chromatin regions (ACRs) as potential cis-regulatory elements. In total, we screened 98 ECRs and ACRs near Onecut1 (Additional File 1, Additional File 2) for their ability to drive reporter activity in the developing retina.

**Figure 1.**
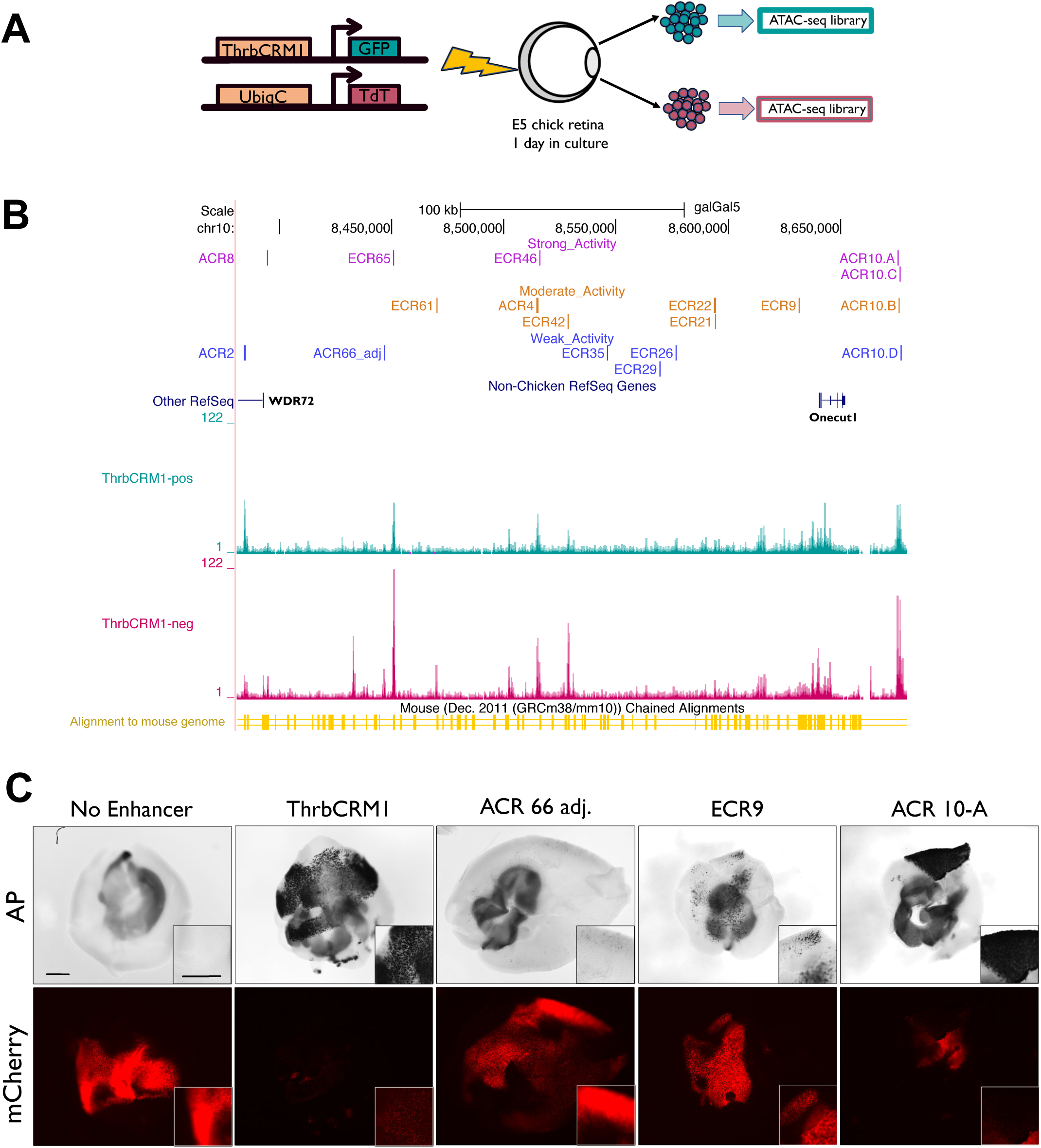
Identification and initial screening for regulatory elements active in E5 chick retinae. (A) Generation of ATAC-seq libraries of ThrbCRM1-positive and ThrbCRM1-negative cell populations. Chick retinae at embryonic day 5 (E5) were electroporated ex vivo with both ThrbCRM1::GFP and UbiqC::TdT plasmids and incubated in culture for 18-22 hours prior to dissociation. Dissociated cells were sorted via FACS into GFP and TdT double-positive cells, and GFP-negative, TdT-positive cells. Each population was processed for ATAC-seq. (B) Visualization of aligned ATAC-seq reads to the galGal5 genome in UCSC Genome Browser in intergenic region between Onecut1 and WDR72 (labelled). Active regulatory elements represented by colored, labelled lines based on activity level from alkaline phosphatase assay (C) Alkaline phosphatase reporter assay to screen for regulatory activity. E5 chick retinae were electroporated with CRE::AP plasmids and CAG::mCherry plasmids. Empty Stagia3 vector (No Enhancer) represents the negative control. Scale bar in the first panel represents 500 µm and applies to all.

The first criterion of our screen was that the non-coding elements should be capable of driving reporter expression at E5 in the chick retina. To this end, we used a sensitive alkaline phosphatase (AP) reporter assay. Each potential cis-regulatory region was amplified from chick genomic DNA and cloned into Stagia3 (Billings et al., 2010), a dual GFP and AP reporter vector. Retinae at E5 (HH26) were electroporated with the reporter construct along with CAG::mCherry as the co-electroporation control. These retinae were cultured for approximately 18-22h and then fixed before the AP stain was developed.

Previously identified Thrb reporters served as positive controls as they are known to be active in the E5 chick retina (Emerson et. al., 2013). The empty Stagia3 vector served as the negative control to demonstrate the baseline levels of transcription when no cis-regulatory element is present. At this time point, the majority of the candidate sequences tested did not drive reporter expression above the baseline level defined by the negative control. The active elements that were initially chosen based on evolutionary conservation were largely found to have open chromatin states in the within the ATAC-Seq datasets. All active cis-regulatory elements (Figure 1C, Additional File 3) were categorized as weak, moderate, or strong (Figure 1B) and further examined for activity in cone/HC RPCs.

### Regulatory Activity within the Cone/HC Restricted RPC population

The second criterion of the screen was specificity to the population of fate-restricted early retinal RPCs that express Onecut1. The ThrbCRM1 element is active in the cone/HC restricted RPC population that expresses Onecut1 and Otx2 (Emerson et al., 2013). It has been previously reported that at 20 hours post-electroporation, 30% of ThrbCRM1(+) cells are in S-phase or G2/M (Buenaventura et al., 2018). Therefore, to determine which active elements drove transcription in the same restricted RPC population, we co-electroporated the CRE::GFP reporter constructs with a ThrbCRM1::AU1 reporter into the chick retina at E5 and cultured overnight for approximately 20 hours. Retinal sections were stained for AU1 and GFP and qualitatively evaluated with regards to the specificity of each active enhancer to the Onecut1(+) restricted RPC population marked by the AU1 reporter (Figure 2, Additional File 4).

**Figure 2.**
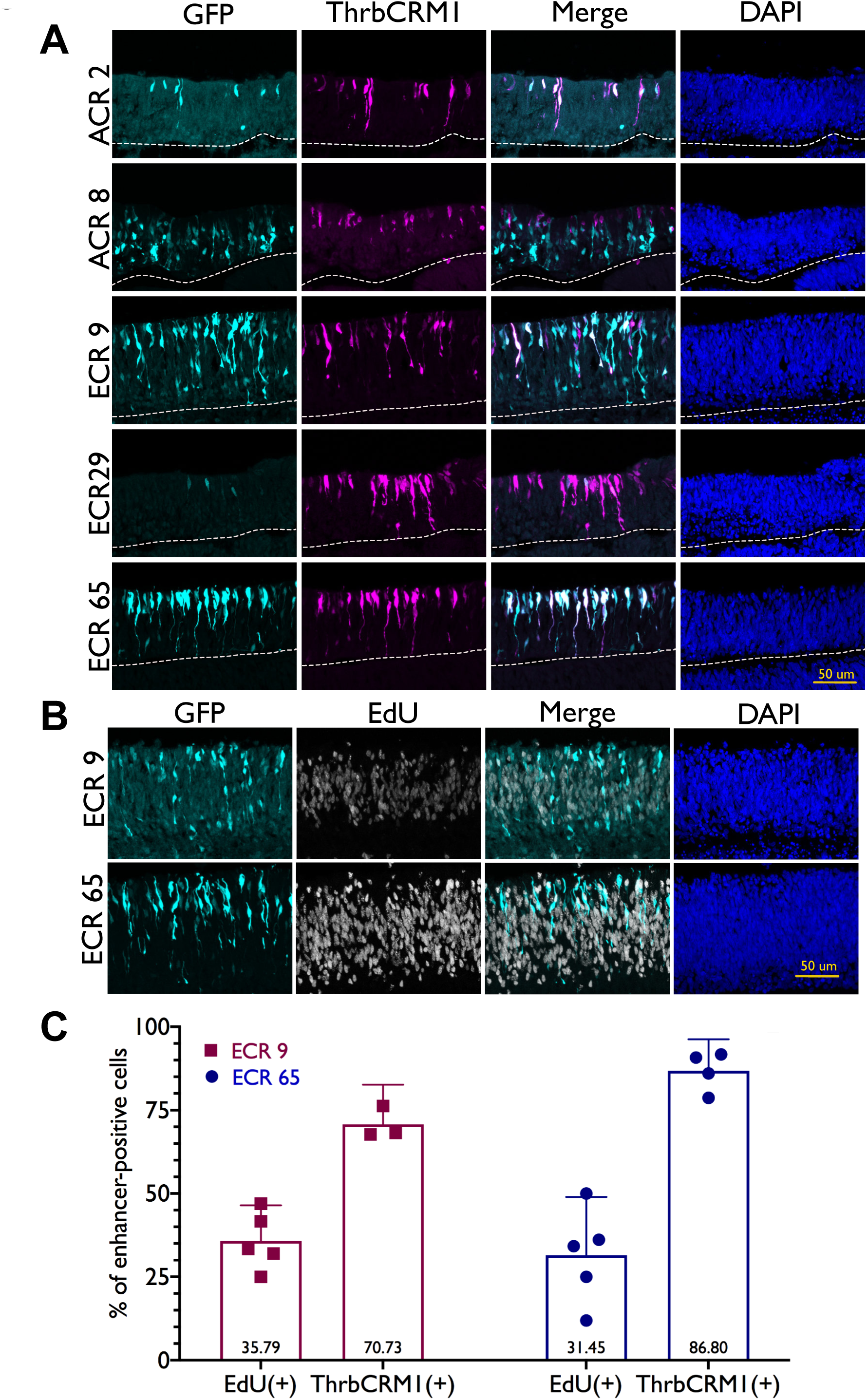
Candidate enhancers ECR9 and ECR65 demonstrate specificity to ThrbCRM1(+) population and overlap with mitotic progenitors. (A) Overlap between CRE::GFP expression (cyan) and ThrbCRM1::AU1 expression (magenta) 18-22 hours after electroporation. DAPI is shown in the last column. Electroporated retina is found above the dotted line (B) Overlap of 1 hous EdU-pulsed cells with ECR9::GFP and ECR65::GFP (cyan) 18-22 hours after electroporation. (C) Quantification of EdU(+) and ThrbCRM1 reporter(+) cells in ECR9::GFP and ECR65::GFP cells. Percentages of EdU(+) cells were calculated from confocal images by determining the number of EdU and enhancer::GFP double-positive cells out of all enhancer::GFP-positive cells. Each point represents a biological replicate with data collected from two images. Percentages of ThrbCRM1(+) cells were calculated from a flow cytometry assay in which retinae were electroporated with ECR9::GFP or ECR65::GFP and ThrbCRM1::TdT and the number of GFP/ TdT double-positive cells of the total GFP(+) population was calculated. Each data point represents a biological replicate. Error bars represent 95% confidence interval.

Many of the active CREs drove reporter expression in both the ThrbCRM1(+) and ThrbCRM1(-) populations, such as ECR42, ECR46 and ACR10-C (Additional File 4). ACR10-B (Additional File 4) activity appears to be distributed throughout the retina with no specific preference for ThrbCRM1(+) cells. Though the populations marked by these CREs overlap with our population of interest, they do not meet the criterion for specificity. Despite driving robust reporter expression in E5 chick retinae, ACR8 and ACR10-A (Figure 2, Additional File 4) appear biased towards ThrbCRM1(-) cells. It is difficult to assess the specificity ECR26, ECR29, and ECR35 to the ThrbCRM1 RPC population due to the weak activity of these enhancers (Additional Files 3 and 4).

ACR2 does not exhibit activity in many cells, but most observed ACR2-positive cells are ThrbCRM1-positive (Figure 2). ECR65 appears to be highly active in ThrbCRM1-positive cells and was qualitatively the most specific enhancer to the ThrbCRM1 population. Likewise, ECR9 activity is highly biased to the ThrbCRM1(+) RPC population but drives reporter activity in some ThrbCRM1(-) cells.

This assay indicates that the active enhancers display a wide range in specificity to the ThrbCRM1(+) cell population (Figure 2A, Additional File 4). ECR 65, ECR 9 and ACR 2 are the most promising candidates for a regulatory element that contains the regulatory information sufficient to promote Onecut1 expression in RPC[CH]s. ECR9 and ECR65 were further assessed for specificity to this restricted RPC cell population as they demonstrated more robust reporter expression than ACR2.

To quantify the specificity of each enhancer to the Onecut1(+) restricted RPCs, E5 chick retinae were electroporated with ECR9:: or ECR65::GFP reporters, ThrbCRM1::AU1, and a co-electroporation control. It was observed that 86.80% of ECR65(+) cells and 70.73% of ECR9(+) cells were also marked by ThrbCRM1 (Figure 2C). However, the ThrbCRM1(+) cell population does not consist of only RPCs. To determine the extent to which each enhancer was active in RPCs, E5 chick retinae electroporated with CRE::GFP constructs were pulsed with EdU for one hour prior to harvest. Approximately 30% of ECR9(+) and ECR65(+) cells are marked by EdU (Figure 2B, 2C), which is comparable to the proportion of EdU(+) cells within the ThrbCRM1 cell population (Buenaventura et al., 2018). Overall, these data provide evidence that the two regulatory elements ECR65 and ECR9 are active in ThrbCRM1 RPCs.

### History of ECR 9 and ECR 65 Activity in Early-born Retinal Cell Types

To further test if these regulatory elements label RPCs which produce cells with the same fates that develop from the ThrbCRM1(+) RPC population, we used a PhiC31 lineage tracing system. In this system, a cis-regulatory element is used to drive the expression of PhiC31, which can activate a GFP responder vector through site-specific recombination to label cells with a history of cis-regulatory activity (Schick et al., 2019). We combined this lineage trace system with immunohistochemistry and cell-specific markers to determine which cell types develop from RPCs marked by ECR65 and ECR9. For comparison, and to demonstrate that not every active regulatory element lineage traces to the same populations at this time point, we also lineage traced ACR2 and ECR42.

It was observed that approximately 40% and 5.5% of ECR9-lineage traced cells were cone photoreceptor (Visinin) and horizontal cell (Lim1) fates, respectively. However, not all ECR9-lineage traced cells correspond to one of these two cells types. Some cells in the ECR9 lineage exhibit long axonal projections which appear to originate from GFP-positive cells in the innermost retina, suggesting that ECR9 is active at some point in the formation of retinal ganglion cells (RGCs). 4.37% of all GFP(+) cells were positive for pan-Brn3, suggesting the presence of RGCs arising from ECR9(+) cells. This may indicate that ECR 9 participates in the gene regulatory network responsible for generating cones, HCs, and RGCs. It is worth nothing that the ThrbCRM1 lineage-trace also includes a similar percentage of pan-Brn3(+) cells as ECR9, despite the lack of inner retinal projections seen with ThrbCRM1 and previous *in ovo* lineage tracing of ThrbCRM1 that detected only a small number of RGCs (Schick et al., 2019). This is potentially due to the antibody’s specificity – it may not mark all Brn3(+) RGC populations or it may be marking cells outside of the target population, including other inner retinal cells such as horizontal or amacrine cells.

Under the same experimental conditions, ECR65::PhiC31 marked overall fewer cells. However, the cell types with a history of ECR65 appeared biased towards the outer retina (Figure 3A). 63.3% of GFP-positive cells were also marked by Visinin (Figure 3B), confirming that ECR65-positive RPCs are capable of giving rise to cone photoreceptors. Surprisingly, very few ECR65-positive cells are marked by Lim1 despite the clear presence of GFP in inner retinal cells (Figure 3A) and reported data that the ThrbCRM1 population gives rise preferentially to Lim1-positive HCs (Schick et al., 2019). To determine whether these cells may be RGCs or Isl1-positive HC, we stained with pan-Brn3 and Isl1. There were few pan-Brn3(+) GFP(+) cells. However, 5.9% of all GFP-positive cells were marked by Isl1. Lineage tracing of ThrbCRM1 yields similar results, with 6.16% of GFP(+) cells also positive for Isl1 (Figure 3B).

**Figure 3.**
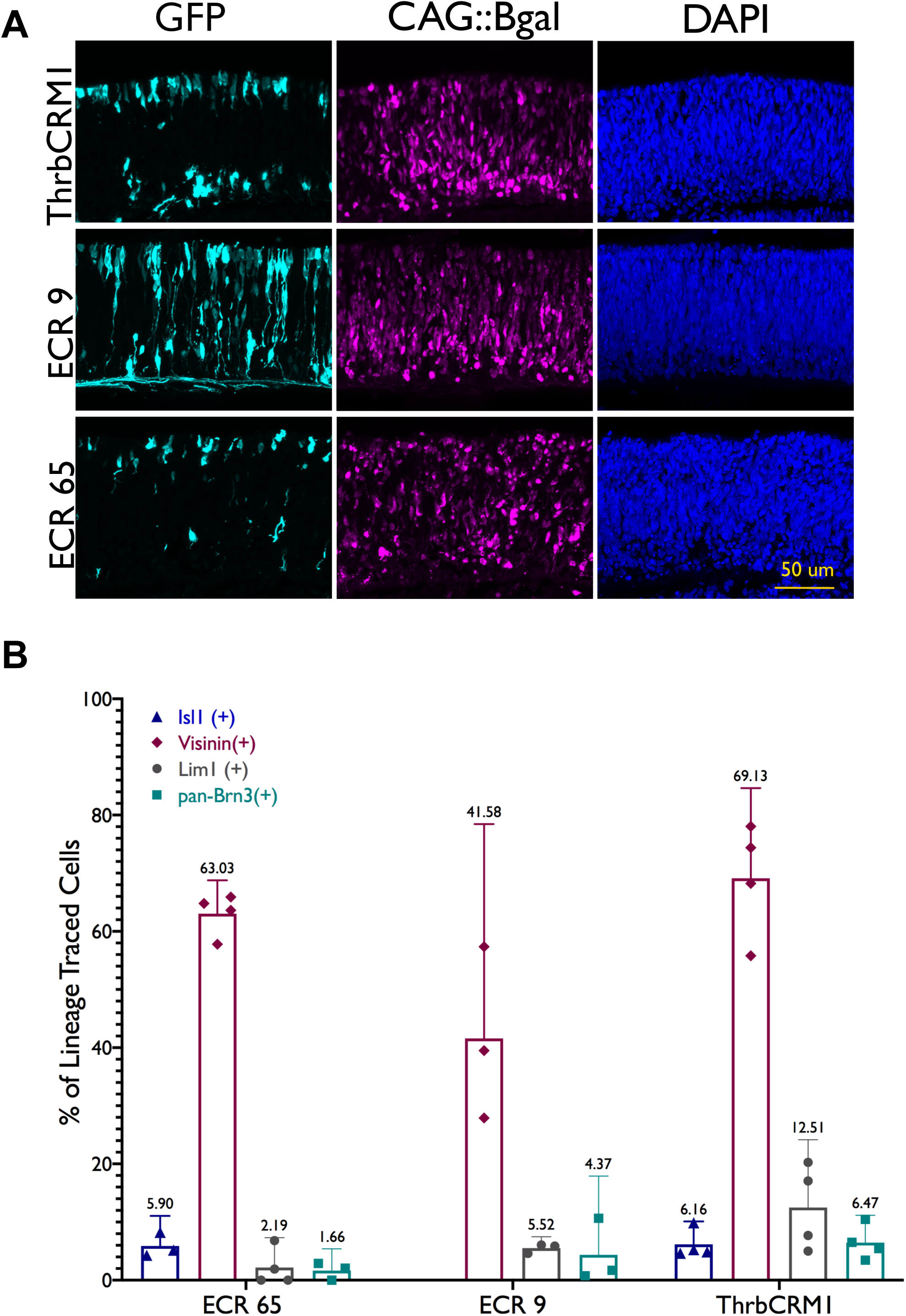
ECR65(+) and ECR9(+) cells lineage trace to similar cell fates as ThrbCRM1(+) population. (A) E5 chick retinas were electroporated with enhancer::PhiC31 constructs and CAG::Bgal as an electroporation control before two days of tissue culture followed by harvest and immunohistochemistry. (A) Retinal sections were stained with GFP, Bgal, and DAPI. (B) Markers of early retinal cell types within lineage traced populations. Sections were stained with the markers Isl1, Visinin, Lim1, and pan-Brn3. Percentages of cells marked by these factors were calculated out of the total number of electroporated Bgal(+) cells per retinal section. Each point represents a biological replicate. Error bars represent 95% confidence interval.

ACR 2 does not lineage trace to very many cells, but is nearly exclusive to the outer retinal cells marked by Visinin, indicating that ACR2’s role in retinal development is specific to photoreceptors, which are predicted to be cones at this timepoint. The lineage tracing of ECR 42 illustrates that this enhancer’s activity is not limited to the cone and horizontal cell fate and marks a much broader RPC population, as evidenced by the pan-retinal distribution of cells with a history of this regulatory element (Additional File 5).

At the conclusion of our screen, eighteen active regulatory elements were identified in the retina, two of which drove GFP reporter expression in a spatial pattern that excluded ThrbCRM1(+) cells. ACR 10.B was found to drive expression non-specifically throughout the retina and three elements, ECR 26; ECR 29; and ECR 35, drove expression too weakly to determine their specificity. ECR 22 was found to mark a population that excludes cone photoreceptors (Gonzalez and Schick et. al., in preparation). ACR2, ECR9, and ECR65 best met the criteria for our screen. These elements are either specific or strongly biased to the ThrbCRM1(+) population, mark early retinal RPC cells expressing Onecut1, and are active in cells that give rise to three early retinal fates: cone photoreceptors; horizontal cells; and RGCs. As the lineage marked by ACR2 is largely specific only to a smaller population of cone photoreceptors, the remainder of this study is focused on evaluation of ECR 9 and ECR 65.

### Bioinformatic Analysis of Regulatory Element Sequences

Though reporter assays indicate that ECR65 and ECR9 drive transcription in the ThrbCRM1(+) population during the cone and HC specification windows in the chick retina, we do not know what role these regulatory elements play in the GRN that gives rise to cone photoreceptors and horizontal cells.

To determine which transcription factors bind to ECR 65 and ECR 9, we first attempted to identify conserved motifs present within the sequence. We used UCSC Blat (Kent, 2002) to find homologous sequences to the originally identified chick ECR 65 sequence from the golden eagle, barn owl, American alligator, thirteen-lined ground squirrel, northern treeshrew, chimpanzee, and human. ECR 65 is well-conserved among all of the avian species as well as the American alligator and is conserved between avians and mammals.

When searching for the 515 bp chick sequence in the mouse genome, UCSC BLAT returned a 214 bp homologous stretch (Additional File 6A). This 214 bp mouse sequence contained a 58 bp stretch that did not align to the chick sequence. In contrast with chick ECR 65, which is wholly located within accessible chromatin, only 150 bp of the homologous mouse sequence are in accessible chromatin region. The accessible chromatin extends past the homologous sequence. Mouse ECR 65 (mECR65), as defined by chromatin accessibility, is 295 bp long (Additional File 6A). In summary, ECR 65 demonstrated both a large sequence divergence and an apparent shift in chromatin accessibility between avian and mammalian species.

To assay this 295 bp mECR65 element for activity, we cloned it into the Stagia3 vector. When mECR65::GFP was electroporated into the chick retina at E5 along with ThrbCRM1::AU1, mECR65(+) cells were observed in the ThrbCRM1 population, which suggests that despite the sequence divergence, mECR65 has retained the regulatory information for activity in these cells. MEME motif analysis (Bailey and Elkan, 1994) of the various species-specific ECR 65 sequences revealed that three motifs appeared conserved between avian and mammalian species (Additional File 6B). It is therefore likely the TF binding sites important for ECR 65 activity are within the conserved Motifs 1, 5, and 2 (Figure 4).

**Figure 4.**
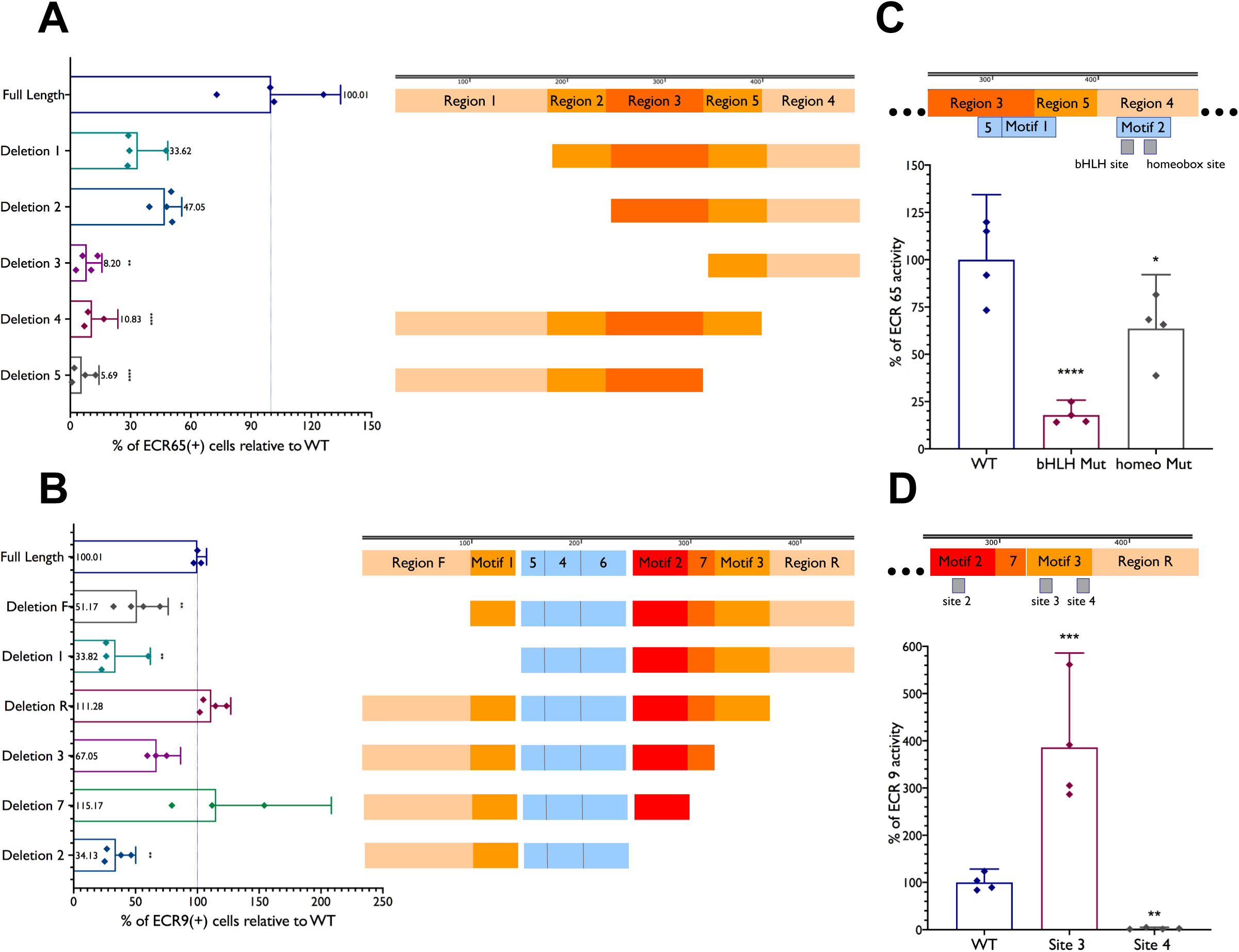
Deletions and mutations reveal sites important for regulatory activity. E5 chick retinae were electroporated with full length (A) ECR65 or (B) ECR9 driving TdT along truncated or mutated versions of the enhancers and CAG::IRFP as a co-electroporation control. Retinae were cultured for 18-22 hours before dissociation and analysis by flow cytometry. Labelled blocks in (A) and (B) represent the enhancer constructs labelled along the Y-axis. Grey blocks in (C) and (D) denote putative TF binding sites. Deleted versions of enhancers were oriented in the expression vector such that the truncated end is farther from the TATA box. Percent of enhancer activity was calculated as a ratio between the total GFP(+) cells and the total TdT(+) cells and scaled to the activity of the full-length enhancer. Error bars represent 95% confidence interval. See Methods section for statistical tests.

The ECR 9 sequence is within accessible chromatin in both chick and mouse early retinal cells. The ECR 9 sequence used was cloned out of the mouse genome and as seen above, is able to drive transcription of a reporter in chick retinal cells during the peak of cone and horizontal cell development. Though the sequence tested is over 400bp long, only a ∼270 bp span is conserved between mouse and chick. Motif analysis of this enhancer through MEME returns seven motifs, of which six are conserved between birds and mammals (Figure 4, Additional Files 6C and 7).

### Deletions and Mutations of Regulatory Element Sequences

To take an unbiased approach to determine which regions of ECR 65 are functional, five serial deletions of the chick ECR 65 sequence were tested for their ability to drive reporter activity (Figure 4, Additional File 7). The full-length/wild type version of ECR65 driving TdTomato was compared against the truncated versions driving GFP using flow cytometry. To ensure that any observed effects were not due to changes in the proximity between TF binding sites and the TATA box, all deletions were oriented such that the distance of all remaining sequence to the TATA box was preserved and these deletion constructs were compared to the full-length enhancer in the same orientation. To determine the effect of each deletion, we calculated the percent of GFP(+) cells relative to the amount of TdT(+) cells. WT ECR65::GFP marks about 56-65% as many cells as WT ECR65::TdT (Additional File 8). When these control values were normalized to 100%, ECR 65 Deletions 1-3 respectively marked 33.62%, 42.05% and 8.2% as many cells as the control WT ECR65::GFP (Figure 4A). When deletions were made on the other end of the regulatory element, GFP driven by ECR65 Deletion 4 or Deletion 5 respectively marks 10.82% and 5.68% as many cells as WT ECR 65. Region 4 also encompasses MEME-predicted Motif 2. The severe loss of GFP expression upon deleting the 80 bp Region 4 suggests that Region 4 of ECR 65 contributes significantly to the activity of this regulatory element.

ECR 65 Motif 2 within Region 4 contains a potential binding site for a bHLH transcription factor. bHLH binding sites, known as E-boxes, typically follow the sequence CANNTG. Mutation of this potential E-box sequence resulted in a significant loss of enhancer activity (Figure 4C, Additional Files 7 and 8). This result suggests that the functional sequence within Region 4 may be this 6-bp motif, predicted by TOMTOM (Gupta et al., 2007) to bind the transcription factor NHLH1/NSCL1. Another mutation within Region 4 encompassing a predicted homeobox TF binding site resulted in some loss of GFP reporter activity (Figure 4C, Additional File 8).

A similar deletion strategy was used to investigate ECR 9 and reporter constructs were tested which removed the ECR9 sequences on either side of the MEME-identified motifs as well as Motif 1, 2, 3, and 7 (Figure 4, Additional File 7). As a control, we calculated the percent of GFP(+) cells relative to the amount of TdT(+) cells marked by full length versions of ECR9 (Additional File 8). Once again, we ensured that deletions were orientated away from the TATA box and compared only to full-length ECR9 of the same orientation. Deletions of Region R, Motif 3, and Motif 7 did not result in a significant change to ECR9 activity. However, deletions of Motif 2, Motif1 and Region F all resulted in a decrease of ECR9 activity as compared to the full-length enhancer. Examination of the sequence for putative TF binding sites led to four predicted bHLH sites (Figure 4, Additional File 7). Site 1 and Site 2 are located in Region F and Motif 2, respectively. Sites 3 and 4 are both located within Motif 3. We hypothesized that if any of these sites were important for ECR9 function, mutation of one or more of them directly would result in a change in reporter expression. Mutation of Site 4, located within Motif 3, resulted in a 97.5% loss of ECR9 activity (Figure 4D, Additional Files 7 and 8).

### Interactions with bHLH Transcription Factors

Our screen for cis-regulatory elements that could regulate Onecut1 expression in early restricted RPCs led to ECR9 and ECR65, which appear to be active in overlapping populations of early RPCs that give rise to distinct subsets of retinal cell types. The tested serial deletions and mutations suggest that both of these elements have critical regulatory input from bHLH family transcription factors. In addition to the bioinformatically predicted Nhlh1 binding site in ECR65, bulk RNA-seq indicated that NeuroD1, NeuroD4, NeuroG2, and Atoh7 transcripts were enriched in the ThrbCRM1(+) population (Buenaventura et. al., 2018) and therefore also candidates to interact with these two cis-regulatory elements. To explore these possibilities, each of the five bHLH factors was overexpressed under the control of the ubiquitous CAG promoter in the E5 chick retina along with ECR65 and ECR9.

Under control conditions in which CAG did not drive any open reading frame, we calculated the percent of ECR9(+) or ECR65(+) cells out of the total electroporated population. Overexpression of either NeuroD1, NeuroG2, and NeuroD4 induced an increase in ECR9 reporter output, while Nhlh1 and Atoh7 did not. Confocal microscopy showed that the increase in ECR9(+) cells upon overexpression NeuroD1, NeuroG2, and NeuroD4 was predominantly in the inner retina (Figure 5B). However, individual overexpression of these three bHLH factors is not sufficient to increase activity of the ECR9 Site 4 mutation (Figure 5D).

**Figure 5.**
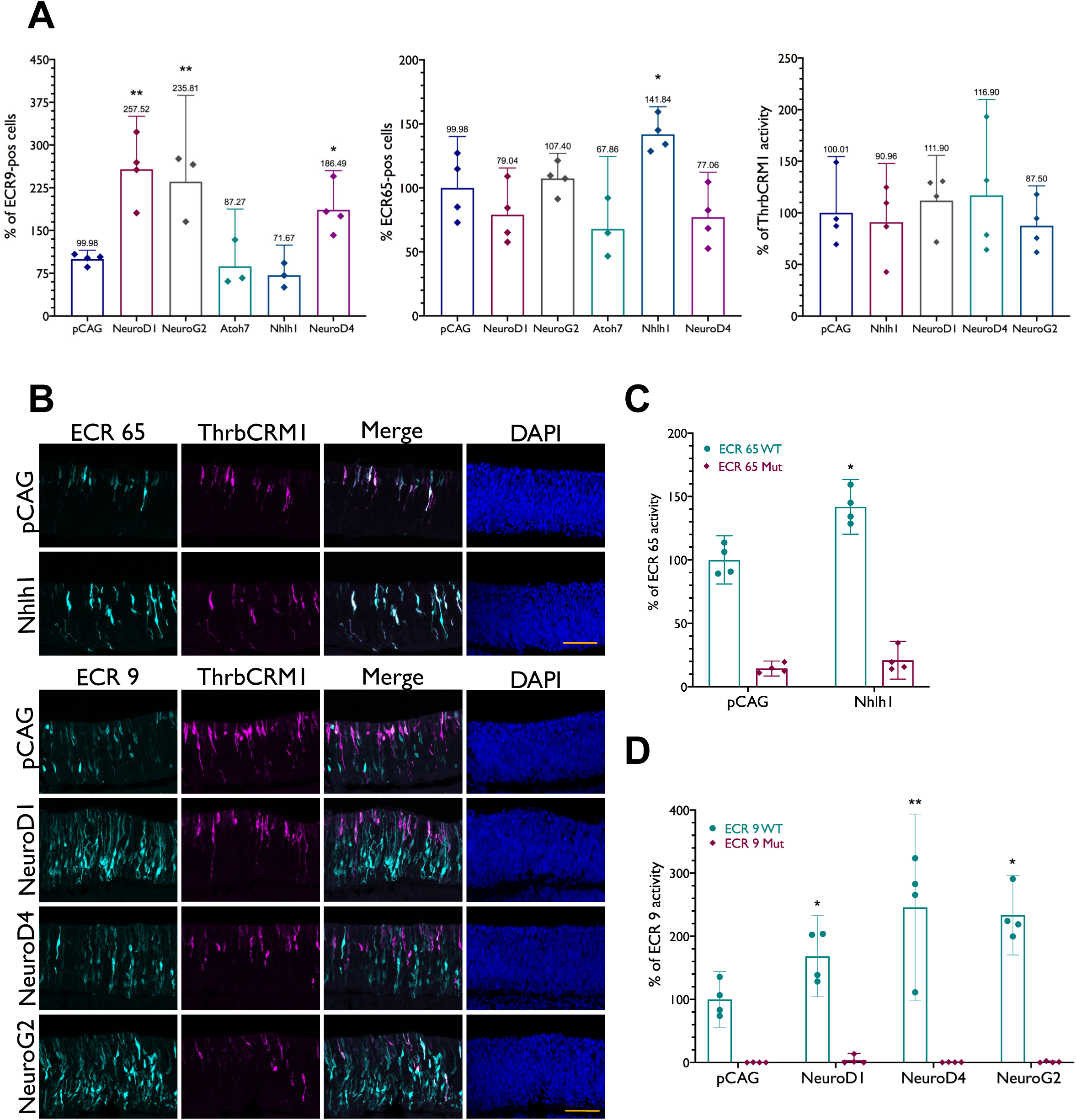
Effect of overexpression of bHLH factors on regulatory activity of ECR9, ECR65, and ThrbCRM1. **(A)** E5 retinae were electroporated with Nhlh1, NeuroD1, NeuroD4, NeuroG2 or Atoh7 under the control of the CAG promoter in combination with ECR65, ECR9 or ThrbCRM1 driving either TdT or GFP. Percentages of ECR9(+), ECR65(+) and ThrbCRM1(+) cells were calculated out of the total cells marked by co-electroporation control CAG::IRFP. **(B)** Retinae were electroporated with ECR9::GFP or ECR65::GFP in combination with ThrbCRM1::AU1 and a CAG::bHLH plasmid. **(C,D)** Comparison of the effect of bHLH overexpression on mutant and WT versions of ECR9 and ECR65. Percentages were calculated out of total number of CAG::IRFP(+) cells. Percentages of WT enhancer(+) cells in the control condition were scaled to 100% for C, D. Percent of enhancer(+) cells in the empty CAG vector control condition were normalized to 100% for A. Error bars represent 95% confidence interval. In all graphs, each datapoint represents a biological replicate. See Methods for statistical tests

We detected no change to the ECR65 reporter output upon misexpression of NeuroD1, NeuroD4, NeuroG2, or Atoh7 (Figure 5A). However, overexpression of Nhlh1 led to a significant increase in the number of GFP(+) cells. Confocal microscopy data showed that the population marked by ECR65 appears to expand towards the inner retina upon Nhlh1 overexpression, with the morphology of some cells resembling multipotent RPCs marked by Vsx2 (Figure 5A, 5B). Furthermore, Nhlh1 overexpression was unable to rescue the loss of GFP activity seen in ECR65bHLH Mut1::GFP, suggesting that the site CATCAG within ECR65 is not only required for regulatory activity, but possibly mediates the interaction between ECR65 and Nhlh1 (Figure 5C).

Despite their effects on ECR9 and ECR65 activity, none of the four candidate bHLH factors were able to increase GFP reporter output driven by ThrbCRM1 (Figure 5A) or drive any changes in the spatial activity of the enhancer in the chick retina at E5 (Figure 5B). In addition, none of the bHLH genes were sufficient to ectopically induce ThrbCRM1 activity in the P0 mouse retina, suggesting that these genes were not sufficient to induce Onecut1 expression at this time (Additional File 9). These results suggest that either ECR9 and ECR65 activation may occur after ThrbCRM1 activation or that the interactions between the bHLH factors and enhancers are not sufficient to induce Onecut1 expression.

### Timeline of Regulatory Element Activity

ECR65::TdT and ECR9::GFP were electroporated with ThrbCRM1::GFP or ThrbCRM1::TdT respectively into retinas that were then cultured for 8 hours to assess whether the two regulatory elements activated at the same time as ThrbCRM1. ECR65, ECR9, and ThrbCRM1 are all able to drive reporter expression 8 hours after electroporation. At this early time point, similar to the 18-22 hour time point, all ECR65(+) cells are ThrbCRM1(+). ECR9 activity at 8 hours is also similar to its activity at 18-22 hours, as there are both ThrbCRM1(+) and ThrbCRM1(-) cells in this population (Figure 6A). Some of the ECR9(+) ThrbCRM1(-) cells in the inner part of the developing retina also appeared to be Vsx2(+) or EdU(+) (Figure 6B, 6C). These results indicate that onset of ECR9 and ECR65 activity is not later than or dependent on ThrbCRM1 activation.

**Figure 6.**
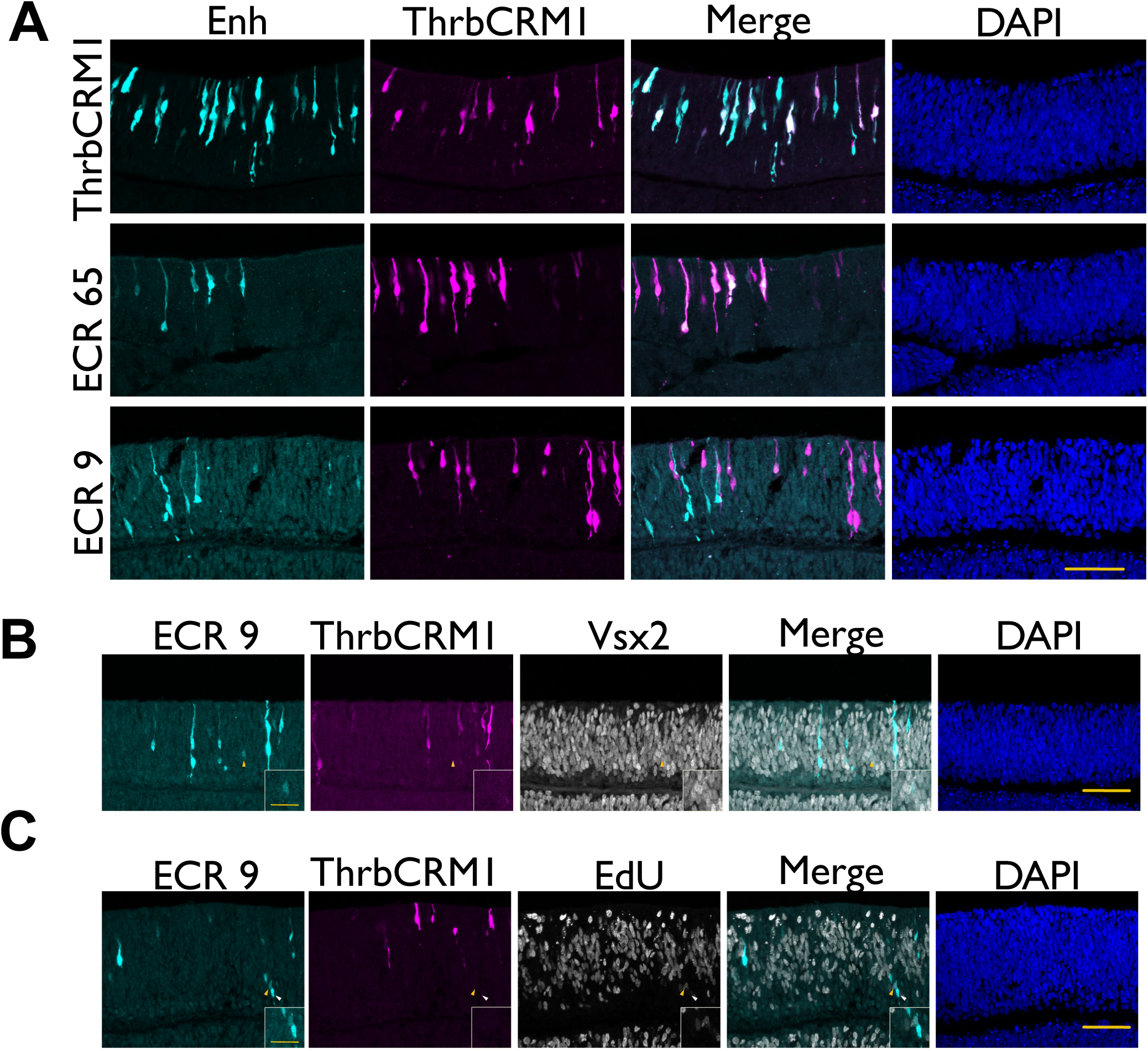
Onset of ECR9 and ECR65 activity compared to ThrbCRM1. (A,B) E5 retinae were electroporated with ThrbCRM1::TdT and either ThrbCRM1::GFP, ECR65::GFP or ECR9::GFP and cultured for 8 hours before harvest. Immunohistochemistry was used to amplify GFP and TdT signal. Yellow arrow in (B) indicates a cell that is ThrbCRM1::AU1(-), ECR9::GFP(+), Vsx2(+). (C) E5 retinae electroporated with ECR9::GFP and ThrbCRM1::TdT were cultured for 8 hours and pulsed with EdU from hour 7-8. Yellow arrow indicates a cell that is ECR9::GFP(+) ThrbCRM1::AU1(-) EdU(+). White arrow indicates a cell that is ECR9::GFP(+) ThrbCRM1::AU1(-) EdU(-). Scale bars in A, B, and C represent 50 µm and apply to all panels. Scale bars in B and C insets represent 20 µm and apply to all panels.

### Molecular Events Upstream of ThrbCRM1 Activity

We then sought to determine whether any of the bHLH factors which impact ECR9 and ECR65 activity affected one or more of the factors upstream of ThrbCRM1 such as Onecut1 and Otx2, or affected expression of the multipotent gene Vsx2. Overexpression of NeuroD1, NeuroD4, or NeuroG2 individually resulted in an increase of electroporated Otx2(+) cells (Figure 7A). Together with the increase in ECR9::GFP(+) cells, this would suggest that the newly GFP(+) cells are expressing Otx2 and indeed there is no change in the proportion of GFP(+) Otx2(+) cells upon overexpression of each bHLH factor (Figure 7A). We then examined whether this enhancer-marked population shared a relationship with the Vsx2(+) multipotent RPC population. Vsx2, shown to be largely absent in the ThrbCRM1(+) restricted RPC population, marks 48.8% of all electroporated cells after one day in culture while ECR9(+) Vsx2(+) comprise less than 1% of all electroporated cells. Overexpression of bHLH factors led to an increase in ECR9(+)Vsx2(+) cells. There was a trend for Vsx2(+) cells within the electroporated population to decrease upon bHLH overexpression but this was not statistically significant (Figure 7A).

**Figure 7.**
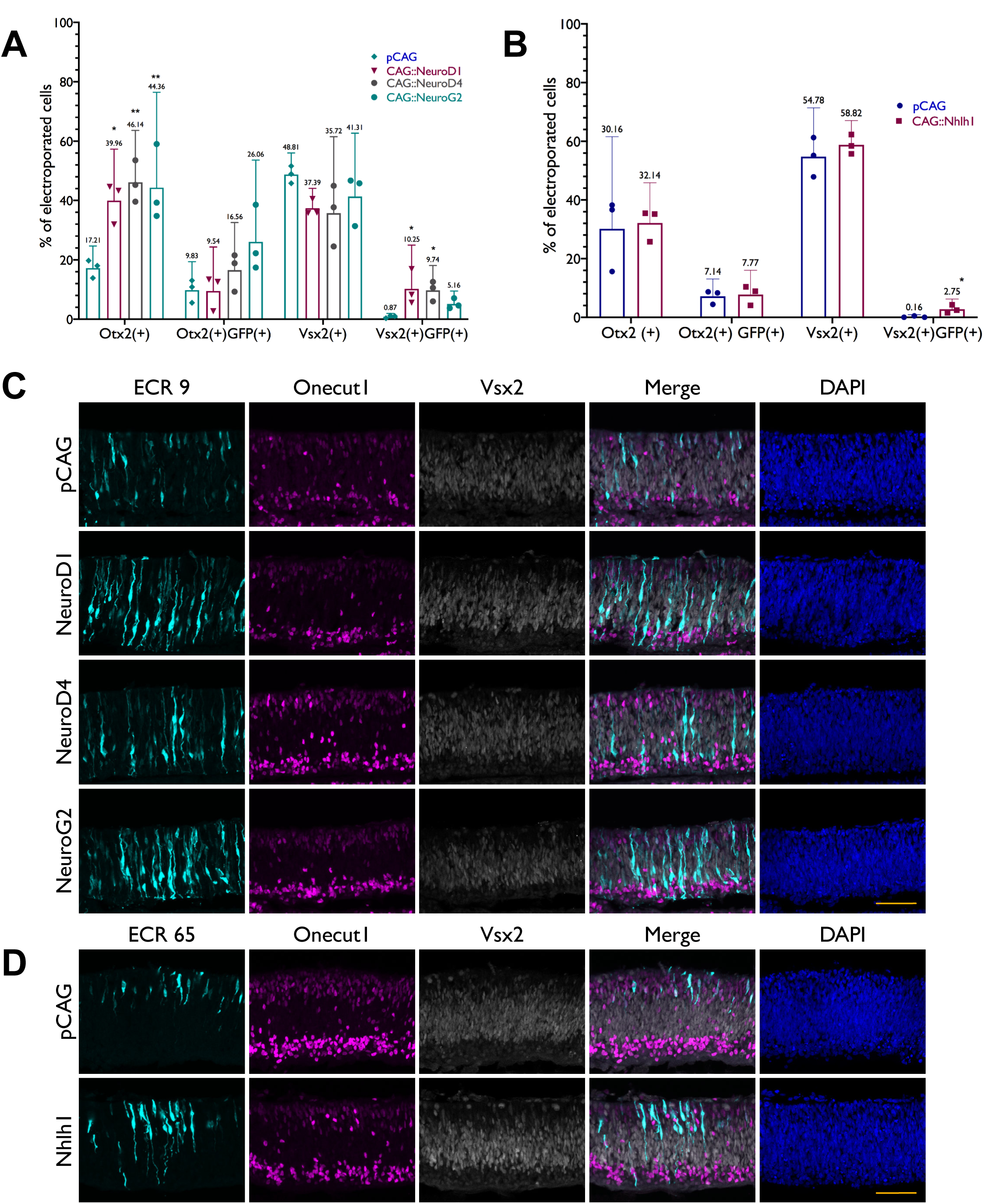
Co-localization of bHLH-induced activity of ECR9 and ECR65 with early retinal development markers. E5 retinas were electroporated with ECR9::GFP or ECR65::GFP and their corresponding bHLH factors and cultured for 18-22 hours before harvest and immunohistochemistry. (A) Cell quantitation derived from confocal images of retinas stained for ECR9::GFP (enhancer), co-electroporation marker Bgal, and either Vsx2 or Otx2. Percentages of Otx2(+) or Vsx2(+) cells as well as GFP(+)Otx2(+) or GFP(+)Vsx2(+) cells were calculated out of the total number of Bgal(+) cells. Error bars represent 95% CI (B) Same as for (A) but for ECR65::GFP. (C) Confocal images of retinas were stained for GFP (enhancer), Onecut1 and Vsx2. Scale bar represents 50 µm.

Overexpression of Nhlh1 does not result in any change to electroporated Otx2(+) cells nor to those cells positive for both Otx2 and ECR65::GFP. Staining with Vsx2 confirmed that there is an increase in ECR65(+) Vsx2(+) cells upon overexpression of Nhlh1 (Figure 7B), as suggested by the data from Figure 5, though the overall percent of Vsx2(+) cells out of the total electroporated population does not change. This result may indicate that Nhlh1 overexpression induces some ECR65 activity within the Vsx2(+) cell population.

Onecut1 staining was used to qualitatively assess the relationship between the enhancers ECR65 and ECR9, the four candidate bHLH factors, Vsx2, and Onecut1. ECR9(+) Onecut1(+) cells appear biased towards the outer retina, where ECR9(+) ThrbCRM1(+) cells are also observed (Figure 2A), whereas ECR9(+) Vsx2(+) cells are more common towards the inner retina (Figure 7C). This does not change when NeuroD1, NeuroD4, or NeuroG2 are overexpressed. However, it appears that many of the strongly GFP-pos cells that appear in the inner retina in response to bHLH overexpression are neither Onecut1(+) nor Vsx2(+).

We observed that under control conditions ECR65 activity overlaps greatly with Onecut1 expression and there are very few ECR65(+) cells which express both OC1 and Vsx2 (Figure 7D). However, ECR65(+) cells which express Vsx2 but not Onecut1 appeared more frequently upon Nhlh1 overexpression.

### Relationship Between bHLH Factors and Early-born Retinal Cell types

We investigated the possibility of these factors affecting retinal cell types developing from E5 to E7. Previously it has been shown that overexpression of Onecut1 in the postnatal mouse retina can induce an increase in cones and horizontal cells while suppressing the rod photoreceptor fate (Emerson et al., 2013). Here, we overexpressed the four candidate bHLH factors, cultured for two days to mirror the lineage tracing experiments, and stained for the same cell-type specific markers (Additional File 10) to determine whether these bHLHs play a role in the development of the cell types marked by the lineage tracing of ECR65 and ECR9.The lineage tracing of ECR9 in conjunction with the data from overexpression of NeuroD1, NeuroD4, and NeuroG2 suggests that these one or more of these transcription factors may play a role in the development of Lim1(+) horizontal cells or RGCs. However, no conclusive increase of these cell types was observed upon bHLH overexpression. Consistent with previously reported data (Li et al., 1999) overexpression of Nhlh1 was not sufficient to induce an increase in Isl1(+) cells. Lastly, none of the four candidate bHLH factors had an effect on photoreceptors marked by Visinin.

## Discussion

The work presented here is intended to serve as a step towards understanding the distinction between cell fate multipotency and restriction. The expression of multipotent RPC genes such as Vsx2 and genes such as Onecut1 that mark the ThrbCRM1(+) fate-restricted RPC population appear to be mutually exclusive (Buenaventura et al., 2018). It is not understood what gene regulatory networks are involved in the establishment of these two populations. We therefore sought to identify cis-regulatory elements specific to the RPC[CH] population that function upstream of Onecut1 and may be involved in the restriction process.

Our large-scale, multi-step screen for regulatory elements resulted in the identification of three regulatory elements that drove a spatial expression pattern biased to the cone/HC restricted RPC population. In addition to ECR9 and ECR65, we found multiple regulatory elements which drove expression in non-specific expression patterns and two which completely excluded the population of interest. These two regulatory elements, ACR8 and ACR10-A, may be involved in generation or maintenance of the multipotent RPC population that gives rise to a broader range of mature cell types as well as the restricted RPC populations. It may also be that the more broadly-acting or ThrbCRM1(-) elements from this screen require further genomic context to act specifically in Onecut1(+) restricted RPCs. Our reporter assay is demonstrably effective in finding minimal elements that can drive specific spatiotemporal expression patterns, but we cannot be certain that we replicate the genomic function of every assayed element given the importance of chromatin state and the surrounding sequences. Our assay does capture some aspects of genomic context as every active element is found within accessible chromatin regardless of sequence conservation. Highly conserved elements located within closed chromatin in E5 chick retinal cells were not found to be active. Despite this, differential chromatin accessibility may not be a strong indicator of enhancer specificity in cell populations at the same developmental time point. ECR65 in particular is accessible in both ThrbCRM1(+) and ThrbCRM1(-) cells, yet under normal conditions is only active in the ThrbCRM1(+) population, demonstrating that appropriate combinations of transcription factors are still required to confer specific activity to regulatory elements.

The regulatory elements ECR9 and ECR65 mark distinct but overlapping populations of developing chick retinal cells from E5-E6, which includes RPCs (Figure 2). ECR65 has also been identified through DNaseI hypersensitivity in the mouse retina as “OC1 A” (Nadadur et al., 2019) but has not been further characterized. Here, we report that the ECR65(+) population overlaps almost entirely with the ThrbCRM1(+) population of cone/HC RPCs (Figure 2) and excludes the Vsx2(+) multipotent RPC population (Figure 7). By combining bioinformatic analyses of the active enhancer sequences with functional tests and previously published RNA-seq data, we were able to connect the activity of ECR9 and ECR65 to bHLH transcription factors.

The overexpression assays demonstrate specificity in the interactions between the four candidate bHLH factors and the two candidate enhancers. The involvement of these transcription factors in retinal development and specifically in retinal cell fate choice has also been well-documented (Cepko, 1999; Hatakeyama and Kageyama, 2004; Dennis et al., 2019). For example, Nhlh1 RNA is known to be present in Isl1(+) RGCs (Li et al., 1999) and has been observed in scRNA-seq from chick retinal cells to mark a cone photoreceptor subtype (Ghinia-Tegla et al., 2019). When overexpressed alongside both candidate enhancers, Nhlh1 is only able to affect the activity of ECR65 (Figure 5A, 5B) which lineage traces to an Isl1(+) population (Figure 3). However, further investigation is required to identify and characterize Nhlh1(+) developing cone photoreceptors and determine whether they develop from the ECR65(+) population. Similarly, NeuroG2 has been found to be important for RGC genesis (Hufnagel et al., 2010; Maurer et al., 2018) and involved in HC fate choice (Akagi et al., 2004). This role may underlie its interaction with ECR9 (Figure 5A, 5B), which lineage traces to horizontal cells and morphologically characteristic RGCs (Figure 3).

The ECR9(+) population also distinguishes itself with a subpopulation that does not overlap with ThrbCRM1 activity. However, this ECR9 (+) ThrbCRM1(-) population that includes RPCs also does not overlap strongly with the multipotent Vsx2(+) population(Figure 6, Figure 7). Though NeuroD1, NeuroG2, and NeuroD4 were considered candidate TFs largely due to their enrichment in the ThrbCRM1(+) population (Buenaventura et. al., 2018), our data did not indicate that overexpression of individual candidate bHLH factors affected ThrbCRM1 activity on the timescale that we examined. Both the quantitative and qualitative assessment along with the visible increase in both Otx2(+) and ECR9(+) cells in the inner retina suggest that NeuroD1, NeuroD4, and/or NeuroG2 mediate ECR9 activity in a population of Otx2(+) cells distinct from the ThrbCRM1(+) restricted RPC population.

It has been hypothesized that an intermediate restricted RPC, marked by ThrbICR, gives rise to ThrbCRM1(+) restricted RPCs (Schick et al., 2019). Lineage tracing of ThrbICR, which is bound by NeuroD1 (Liu et al., 2008), shows that the cells marked by that element can give rise to RGCs as well as cones and horizontal cells (Schick et al., 2019). In conjunction with the ECR9 lineage tracing data and NeuroD1 overexpression data, ECR9 activity in cells which are neither ThrbCRM1(+) nor Vsx2(+) suggests that ECR9 could also label this intermediate population, referred to as RPCs[CHG] (Figure 8A).

**Figure 8.**
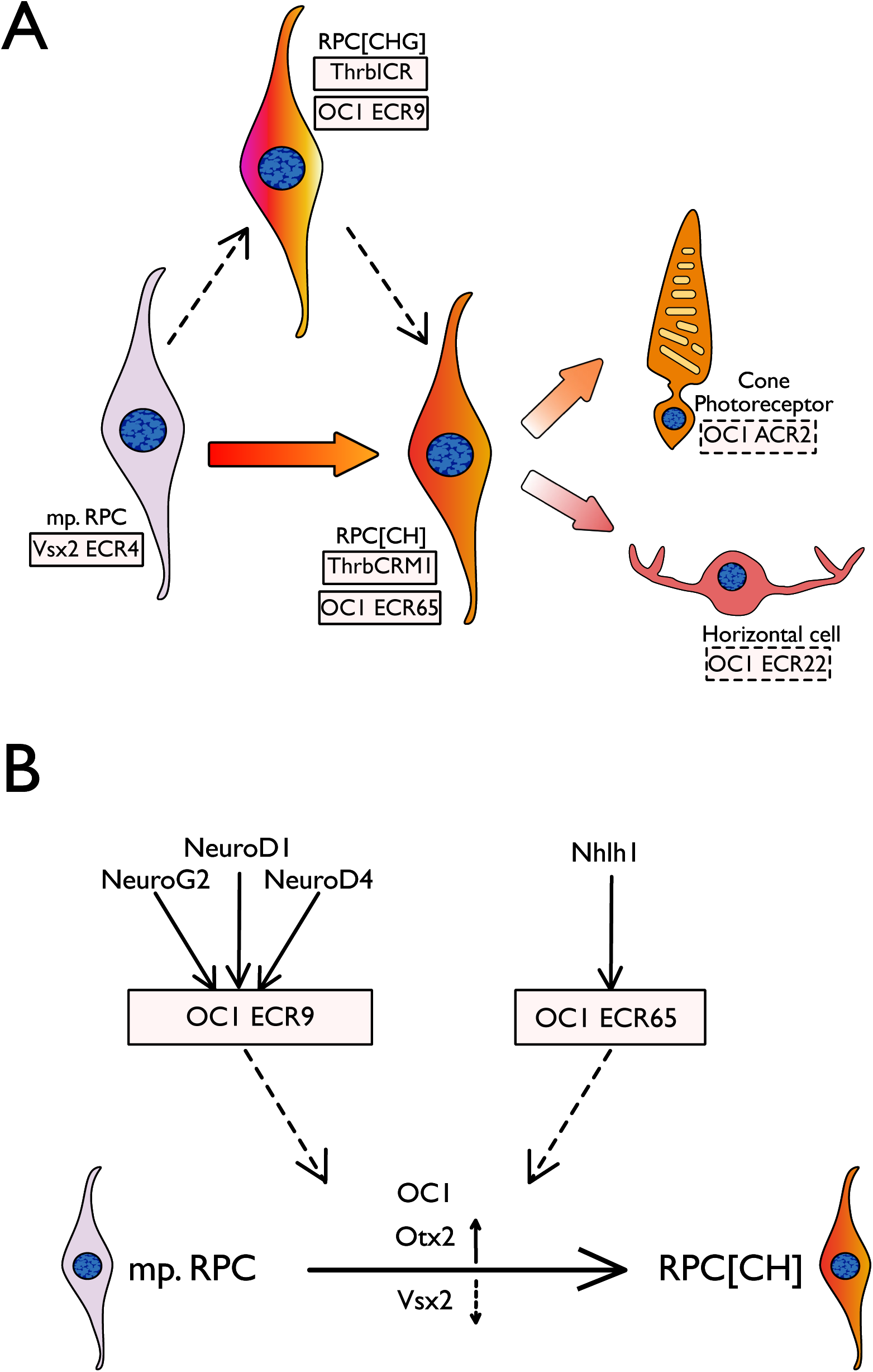
Model of ECR9 and ECR65 roles in cone/HC regulatory network. (A) Cell populations with ECR9 and ECR65 activity in relation to populations marked by previously published elements and OC1-associated elements reported here. Vsx2 ECR4 is active in the multipotent RPC population, whereas OC1 ECR9, OC1 ECR65, ThrbCRM1, and ThrbICR are all active in fate-restricted RPC populations. (B) Molecular events upstream and downstream of ECR9 and ECR65 activity. Multipotent RPCs give rise ultimately to RPC[CH]s, which corresponds to a down-regulation of Vsx2 and an upregulation of OC1 and Otx2. The bHLH factors that are sufficient to activate ECR9 and ECR65 reporter expression are shown.

The regulatory elements uncovered in this and previous screens can therefore be used to mark distinct developing populations in the early vertebrate retina. OC1 ECR65 and ThrbCRM1 are both active in the restricted RPC[CH], while OC1 ECR9 and ThrbICR activity may be marking the hypothesized RPC[CHG] which is distinct from multipotent RPCs characterized by the activity of VSX2 ECR4 (Buenaventura et al., 2018). Other elements found in the screen, such as ACR2 (Figure 2, Additional File 5) and ECR22 (Gonzalez and Schick et. al., in preparation), may be more specific to either the cone photoreceptor precursors or horizontal cell precursors, respectively, from the ThrbCRM1 lineage (Figure 8A). Though ECR9(+) and ECR65(+) cell populations overlap with each other and other CREs such as ThrbCRM1 and potentially ThrbICR, the two enhancers characterized here are unique in their responses to the bHLH factors described above. We hypothesize that the interactions between ECR9 and ECR65 and their respective bHLHs play a role in the transition from multipotent RPCs to the fate-restricted RPC[CH] population (Figure 8B). It may be that one or more of the bHLH factors shown to interact with ECR9 may play a role in coordinating Vsx2 downregulation and activation of Onecut1 expression and ThrbCRM1 activity (Figure 8B).

While we have not determined which factor(s) among NeuroD1, NeuroD4 and NeuroG2 interact with ECR9 *in vivo*, the ability of all three factors to increase ECR9 activity is consistent with a previous study demonstrating their functional redundancy during retinogenesis (Akagi et al., 2004). As ECR9 has multiple putative E-box sites, it may be that some combination of two or all three factors is required for the correct regulatory output of this enhancer. Of the ECR9 bHLH sites, Sites 2-4 all vary in sequence but all three sites and their flanking sequences are highly conserved between mouse, chick and human (Additional File 7). It is also worth noting that while the ECR65 bHLH site shares the same sequence as ECR 9 Site 3, the two enhancers still differ in their ability to respond to Nhlh1 overexpression. This may be due to the differences in their flanking sequences. Previous work has shown that some bHLH factors are able to utilize each other’s binding sites (Mao et al., 2013) and that the sequence flanking the core E-box may also be important for binding affinity (Gordân et al., 2013).

Our results do not suggest that any of these bHLH factors individually are capable of inducing or suppressing any of the early-born cell fates in the chick retina. Previous studies using Xenopus were able to yield an increase in photoreceptors upon overexpression of NeuroD1 and NeuroD4 (Wang and Harris, 2005). In chick, NeuroD1 has been reported to induce more photoreceptors when virally overexpressed in the chick retina at E2 and cultured until E8.5/E9 (Yan and Wang, 1998). Our lack of a similar result may be due to differences in the experimental timepoints, as our experimental conditions included a maximum culture time of 48 hours. Though we were also not able to induce horizontal cell or RGC fates, it may be because the overexpressed factors require the co-expression of other TFs in order to specify or induce particular cell fates. For instance, the prediction of a homeobox TF binding site so close to the bHLH binding site in ECR65 may suggest that both Nhlh1 and a homeobox factor are required to induce the cell fates observed from lineage tracing ECR65. Future studies are required in which bHLH factors are overexpressed in combination with each other and with homeobox factors to determine which factors are sufficient to drive early retinal cell fates in chick.

### Conclusions

This study examined the upstream regulatory events of the Onecut1 gene that occur in chick RPCs. Guided by both sequence conservation and chromatin accessibility, we identified two regulatory elements near the Onecut1 gene, ECR9 and ECR65, that are preferentially active in Onecut1-expressing ThrbCRM1(+) RPCs. We find that both of these elements are predicted to contain bHLH transcription factor binding sites, which are required for activity of these elements. Overexpression of specific bHLH members leads to ectopic activity of ECR9 and ECR65 in Vsx2(+) RPCs. These bHLH factors are able to upregulate endogenous Otx2 expression, a protein normally expressed in fate-restricted RPCs. Taken together, these results suggest a role for bHLH factors in promoting the formation of fate-restricted RPCs from multipotent RPCs through the activation of Onecut1 and Otx2.

## Abbreviations

GRN: gene regulatory network
RPC: retinal progenitor cell
HC: horizontal cell
RPC[CH]: restricted retinal progenitor cell that gives rise to cones, horizontal cells
RPC[CHG]: restricted retinal progenitor cell that gives rise to cones, horizontal cells, retinal ganglion cells
CRE: cis-regulatory element
TF: transcription factor
GFP: green fluorescent protein
TdT: TdTomato

## Declarations

### Acknowledgements

Diego Buenaventura provided guidance with the bioinformatic analyses and Sean McCaffery generated the lineage tracing plasmids for ECR9 and ECR42. Miruna Ghinia-Tegla, Estie Schick, Xueqing Chen and the rest of the Emerson Lab provided valuable feedback. The Lim1 and Isl1 antibodies developed by TM Jessell and S. Brenner-Morton and the Visinin antibody developed by C. Cepko and S. Bruhn were obtained from the Developmental Studies Hybridoma Bank, created by the NICHD of the NIH and maintained at The University of Iowa, Department of Biology, Iowa City, IA 52242. Jeffrey Walker and Jorge Morales provided assistance with flow cytometry and confocal microscopy, respectively.

### Availability of Data and Materials

ATAC-seq data generated from chick retinal cells has been submitted to GEO as data set XXXXXX and will be available upon publication. ATAC-seq data generated from mouse retinal cells is not currently available as it is part of another study but can be made available upon request.

### Authors Contributions

ME and SP generated the ATAC-seq libraries, planned and conducted experiments and wrote the manuscript. NJC led the initial screen shown in Figure 1 and Additional Files 1 and 2. AG, SG, BW, and BS generated plasmids and processed tissue for Figures 1 and 2 and Additional Files 1 and 2. BS also generated mutagenized plasmids for Figure 4. SS generated plasmids and processed tissue for Figures 4 and 5.

### Ethics approval and consent to participate

The City College of New York Institutional Animal Care and Use Committee approved all animal procedures under protocol 932.

### Consent for publication

Not applicable

## Methods

### Animals

Fertilized chick eggs were acquired from Charles River and stored at 16 °C room for a maximum of ten days. Embryonic days were counted from E0 when eggs were moved to a 38 °C humidified incubator for five days.

### ATAC-seq

ATAC-seq libraries were collected and amplified as outlined by Buenrostro et al., 2013. Mouse retinas were collected from embryonic day 12.5 (plug morning equal to time 0.5) embryos, and dissociated using manual douncing. Libraries were analyzed for quality control on Bioanalyzer and Qubit and then sequenced at a depth of 37.5 million reads per sample. Sequenced libraries were prepared for analysis using FASTQGroomer (Blankenberg et al., 2010) on default settings and then analyzed using Bowtie for Illumina (Langmead et al., 2009) with default settings except -X 2000 and – m 1, through the usegalaxy.org (Afgan et al., 2018) web platform. Resulting SAM files were converted to the BAM format (Li et al., 2009) and the BigWig (Kent et al., 2010) format.

### Electroporation

Retinae were electroporated ex vivo as previously described (Schick et al., 2019). CAG::reporter plasmids were used at a concentration of 0.1ug/uL with the exception of CAG::mCherry, which was used at a concentration of 0.04 ug/uL. Reporter plasmids under the control of a tissue-specific enhancer (ex: ThrbCRM1, ECR9, ACR2, etc) were used at a concentration of 0.16 ug/uL For lineage tracing experiments, all plasmids (recombinase, responder plasmid, co-electroporation control) were used at a concentration of 0.1 ug/uL. Recombinase and responder plasmids are described in Schick et. al., 2019. Plasmids used in initial identification screen for cis-regulatory elements used miniprep scale DNA purification (Zymo Research, D4020 or 5 prime/Eppendorf, FastPlasmid kit) and all subsequent experiments used midiprep scale DNA purification (Qiagen, 12145 or 12243).

### Alkaline Phosphatase Staining

Retinas were harvested from culture and fixed in 4% paraformaldehyde (PFA), washed 3X in PBS, and incubated in 1 mL NTM (pH 9.5) buffer for 15 minutes while shaking at a low speed before addition of 1mL NTM with NBT(0.25 mg/mL) and BCIP (0.125 mg/mL). Retinae were incubated with the AP substrates in the dark for 2-3 hours until the positive control was well-stained.

### Immunohistochemistry

Retinas were harvested from culture, prepared for cryosectioning, and 20 micron vertical sections were collected as outlined in Schick et al., 2019. Sections were incubated for 10 minutes in 0.1% Tween (VWR, 97062-332) in PBS (PBT) and then blocked for 1 hour in 5% serum (Jackson ImmunoResearch, Donkey - 017-000121, Goat - 005-000-121) in PBT at room temperature prior to incubation with primary antibodies overnight at 4 C. Sections were washed 3X with PBT, prior to blocking at room temperature for 30 min and then incubated with secondary antibodies overnight at 4 C. DAPI was added at 1 ug/uL in PBT while washing off the secondary antibodies. Sections were then mounted using Flouromount-G (Southern Biotech, 0100-01) and coverslips (VWR, 48393-106). Primary antibodies are listed in the table below. All secondary antibodies were obtained from Jackson ImmunoResearch and suitable for multiple labelling. All Alexa-conjugated secondary antibodies were used at dilution of 1:400 and Cy3-conjugated secondary antibodies were used at a dilution of 1:250 from secondary antibody stocks in 50% glycerol.

| Antibody; dilution | Vendor | Catalog number |
| --- | --- | --- |
| Chick anti-GFP 1:2000 | Abcam | ab13970 |
| Rabbit anti-GFP 1:1000 | Invitrogen | A6455 |
| Mouse anti-AU1 1:2000 | Enzo | ENZ-ABS135-0200 |
| Chick anti-Bgal 1:1000 | Abcam | ab9361 |
| Rabbit anti-RFP 1:250 | Rockland | 600-401-379 |
| Mouse anti-Onecut1 1:200 | Santa Cruz | Sc-376308 |
| Goat anti-Otx2 1:500 | R&D | AF1979 |
| Sheep anti-Vsx2 1:200 | Ex Alpha | x1180p |
| Mouse anti-Visinin 1:250 | DHSB | 764-s |
| Mouse anti-Lim1 1:15 | DHSB | 4F2-C |
| Mouse anti-Isl1 1:50 | DHSB | 39.3F7 |
| Mouse anti-panBrn3 1:100 | Santa Cruz | Sc-390781 |

### Microscopy and Cell Counting

Images of whole AP-stained retinas were acquired using a Zeiss Axiozoom V16 microscope with a 1X objective and Zen 2 Blue 2011 software. All confocal microscopy images were acquired using a Zeiss LSM 710 inverted confocal microscope with a 40x oil immersion objective, 488nm laser, 561 nm laser, 633 nm laser, 405 nm laser, and Zen Black 2015 21 SP2 software at a resolution of 1024 x 1024, acquisition speed of 6 and averaging number of 2. For all confocal images shown, Z-stacks consisting of 8-12 Z planes were collected and are shown as maximum intensity projections. Cells were counted in the Fiji (Schindelin et al., 2012) distribution of ImageJ, using the Cell Counter plug-in developed by Kurt De Vos. In the lineage-tracing experiments and for the EdU assay, all GFP(+) cells which co-localized with DAPI were counted first and cells positive for EdU or for cell-specific markers such as Visinin were counted among this population. In both instances, multiple images per retina were analyzed if necessary to reach at least 45-50 GFP(+) cells. Brightness and contrast were adjusted uniformly across each image in Affinity Designer vector editor (Serif [Europe] Ltd).

### EdU Pulse and Detection

5 uL of 10mM EdU solution was added to 1 mL of culture media during the last hour of incubation before retinas are harvested as described above. EdU was detected using the Click-iT Plus EdU Kit for Imaging (Invitrogen, C10640). The tissue was first incubated for 15 minutes in 0.1% Tween in PBS at room temperature before incubating with the EdU Reaction Cocktail for 30 min in the dark at room temperature. The EdU Reaction Cocktail was then removed with 3 washes of PBT prior to antibody staining.

### Deletions and Mutagenesis

Deletions or truncated versions of regulatory element sequences were generated through the use of PCR primers that began internally within the sequence and excluded the portions of the sequence to be deleted. Mutant regulatory element sequences were generated using overlap extension PCR, in which primers included short sequence mismatches at potential TF binding sites. A second set of primers encompassed the ends of the regulatory element sequence and the restriction sites within the plasmid template for easy cloning of the mutant sequence into the Stagia3 reporter vector. Site 3 and Site 4 within ECR9 were mutated to the sequences “TTAGAC” and “AGGCCA”. The bHLH site within ECR65 Region 4 was also mutated twice, to the sequence “CTGATGAATGGCG” to include the E-box site, 4 bp upstream and 3 bp downstream and to the sequence “TTTCCCAAAG” to include the E-box site and 4 bp upstream. (Figure 4, Additional File 8).

### MEME-suite

To identify conserved motifs within ECR65, we used MEME with settings to find a maximum of 12 motifs, with a width of 6-50 bp each and 2-9 sites per motif. For ECR9, we used settings to find a maximum of 7 motifs, with a width of 6-50 bp each and 2-8 sites per motif. The output from MEME was then used as input for TOMTOM under default settings.

### Multiple Sequence Alignments

The alignments between the chick, mouse, and human sequences for ECR9 and ECR65 (Additional File 7) were produced in Clustal Omega (Madeira et al., 2019) version 1.2.4 with default settings.

### Plasmids

Plasmids containing coding sequences of candidate TF genes were obtained from Transomics. Mouse Nhlh1 (Clone ID: BC051018) and mouse NeuroD4 (Clone ID: BC054391) were cloned using EcoR1 to insert into a modified pCAG vector that allows for EcoR1 flanked insert cloning (Emerson et al., 2013) while mouse NeuroD1 (Clone ID: BC018241), human NeuroG2 (Clone ID: BC036847) and human Atoh7 (Clone ID: BC032621) were cloned using a combination of EcoR1 and Not1 (NeuroD1, NeuroG2, Atoh7) into pCAG::EGFP (Matsuda and Cepko, 2004) such that each coding sequence is under the control of the CAG promoter. The following plasmids were previously reported: CAG::OC1, ThrbCRM1::GFP, ThrbCRM1::AU1 plasmids (Emerson et al., 2013); CAG::iRFP (Buenaventura et al., 2018); Bp::PhiC31 lineage tracing and CAaNa::GFP responder plasmids (Schick et al., 2019); UbiC::TdTomato (Rompani and Cepko, 2008); and TdTomato reporter plasmid (Jean-Charles et al., 2018). The CAG::mCherry and CAG::nucBgal plasmids were constructed by Takahiko Matsuda and reported in (Wang et al., 2014) and obtained from the Cepko lab, respectively. The ThrbCRM1::TdTomato plasmid was made by ligating a Not1/EcoR1 fragment from ThrbCRM1::GFP into the TdTomato reporter plasmid (Jean-Charles et al., 2018). Candidate Onecut1 cis-regulatory elements were amplified from chick or mouse genomic DNA with Herculase II polymerase (Agilent, 600677-51), treated for 10-30 minutes with Taq polymerase (Qiagen, 201203) to generate Adenine overhangs, and ligated into PGemTeasy (Promega, A1360). Inserts were sequence verified by Sanger sequencing (Genewiz) and moved into Stagia3 after EcoR1 digestion. In cases where elements contained EcoR1 sites, the original sequences used to amplify candidate elements were used to generate modified oligos with Xho1, Sal1, or Mfe1 restriction sites to allow for PCR-amplification of elements from the verified PGemTeasy clones and subsequent insertion into an appropriately digested Stagia3 plasmid. As candidate elements could be inserted into Stagia3 in two orientations, for some elements both possible orientations were tested.

### Dissociation and Flow Cytometry

Upon harvest, retinae were dissociated into single cells as described in Schick et al, 2019 using papain (Worthington, LS003126) and an activation solution of L-cysteine (VWR, 97063-478) and 10mM EDTA at 37 C. 10% FBS (ThermoFisher, A3160602) solution in DMEM (Life Technologies, 11995-073) was used to stop the dissociation .Cells were further digested with DNaseI (Sigma, 4536282001) and subsequently washed in DMEM prior to fixing in 4% PFA. Dissociated cells were then analyzed on a BD LSRII machine using the 488nm, 561nm, and 633nm lasers. The collected data was analyzed using FlowJo version 10.4.2.

### Statistical tests

Statistical tests were conducted using GraphPad Prism. Data sets were tested for normality (Shapiro-Wilk) prior to ANOVA, Kruskal-Wallis, or t-tests. Significant Kruskal-Wallis or ANOVA p-values were followed up with Dunn’s or Dunnett’s post hoc test, respectively.

| Figure | Exp. Condition | Dunnett p-value | Dunn p-value | Unpaired t-test |
| --- | --- | --- | --- | --- |
| **Figure 4a** | ECR65 Deletion 3 |  | 0.0011 |  |
|  | ECR65 Deletion 4 | < 0.0001 |  |  |
|  | ECR65 Deletion 5 | < 0.0001 |  |  |
| **Figure 4c** | ECR9 Deletion F |  |  | 0.0016 |
|  | ECR9 Deletion 1 |  |  | 0.006 |
|  | ECR9 Deletion 2 |  |  | 0.0025 |
| **Figure 4b** | ECR65 bHLH Mut A | < 0.0001 |  |  |
|  | ECR65 hb Mut | 0.0242 |  |  |
| **Figure 4d** | ECR9 site 3 Mut | 0.0007 |  |  |
|  | ECR9 site 4 Mut | 0.001 |  |  |
| **Figure 5a** | ECR65 + Nhlh1 | 0.046 |  |  |
|  | ECR9 + NeuroD1 | 0.0019 |  |  |
|  | ECR9 + NeuroG2 | 0.0085 |  |  |
|  | ECR9 + NeuroD4 | 0.021 |  |  |
| **Figure 5c** | ECR65 + Nhlh1 | 0.046 |  |  |
| **Figure 5d** | ECR9 + NeuroD1 | 0.0137 |  |  |
|  | ECR9 + NeuroD4 | 0.01 |  |  |
|  | ECR9 + NeuroG2 | 0.0164 |  |  |
| **Figure 7a** | Nhlh1 Vsx2 + GFP |  |  | 0.035 |
|  | ECR9 + NeuroD1 Otx2 | 0.0245 |  |  |
|  | NeuroD1 Vsx2 + GFP | 0.0287 |  |  |
|  | ECR9 + NeuroG2 Otx2 | 0.0098 |  |  |
|  | ECR9 + NeuroD4 Otx2 | 0.0069 |  |  |
|  | NeuroD4 Vsx2 + GFP | 0.037 |  |  |

