## Supplemental Information for "Cis-regulatory analysis of Onecut1 expression in fate-restricted retinal progenitor cells"

**Additional File 1 - Genomic map of potential regulatory elements identified near Onecut1 gene locus.** Coding regions of Onecut1 and WDR72 from multiple species are marked by blue bars. Yellow bars and lines at the bottom indicate sequence conservation between chick and mouse (mm10 assembly) genomes. Chromatin accessibility reads from the ThrbCRM1-positive population are shown in teal, reads from the ThrbCRM1-negative cell population are shown in magenta. Peaks in magenta may be cut off as the data was scaled to optimally visualize the ThrbCRM1 chromatin accessibility. Black bars at the top indicate potential regulatory elements, labelled as Evolutionary Conserved Region (ECR) if originally identified through sequence conservation and Accessible Chromatin Region (ACR) if identified through ATAC-seq data. Elements which could not be cloned or assayed for activity are marked with an asterisk.

**Additional File 2 –** ID and genomic coordinates of all tested sequences in the galGal5 chick genome assembly, with the exception of ECR9 which is in the mm10 mouse genome assembly.

**Additional File 3. Regulatory elements active in E5 chick retinae**. E5 chick retinae electroporated with Enhancer::AP plasmids and CAG:: mCherry plasmids and cultured for 1 day prior to alkaline phosphatase assay. Shown are the AP reporter signal on top and the mCherry signal on bottom. Insets in AP panels show zoomed in areas of reporter activity. Scale bar in last panel represents 500 µm and applies to all.

**Additional File 4. Overlap between ThrbCRM1 activity and activity of eleven candidate enhancers.** E5 chick retinae were electroporated with enhancer::GFP (cyan) constructs as well as ThrbCRM1::AU1 (magenta) constructs and cultured for 18-22 hours prior to antibody staining with GFP, AU1 and DAPI (nuclei) to determine which enhancers marked the same cell population as ThrbCRM1. Scale bar in last panel represents 50 μm and applies to all.

**Additional File 5. Lineage tracing of regulatory elements reveals range in specificity.** ACR2::PhiC31 and ECR42::PhiC31 were electroporated into E5 chick retinae with a PhiC31 GFP responder plasmid and CAG::Bgal and cultured for two days before harvest and staining with GFP to label cells with a history of PhiC31 expression and Bgal to label all electroporated cells. Scale bar in top right panel represents 50 μm and applies to all.

**Additional File 6. Conservation of sequence, chromatin state and function of ECR65.** (A) The entirety of chick ECR65 (purple bar) aligns to open chromatin in the chick genome. The homologous mouse sequence (grey bar with red lines) only partly aligns to the open chromatin region in the mouse. Mouse ECR65 (long black bar) is a longer region of open chromatin. Motifs 1, 5, and 2 (small labelled black bars) are conserved between both Mouse ECR65 and Chick ECR65. Insets show zoomed out genomic area to include surrounding closed chromatin. (B) Mouse ECR65::GFP was electroporated into E5 chick retina along with ThrbCRM1::AU1 and cultured for 18-22 hours before harvest and immunohistochemistry. Retinae were stained for GFP, AU1 and DAPI to examine overlap between GFP and AU1. Scale bar shown in last panel represents 50 μm and applies to all. (C) Chromatin accessibility at the ECR9 region in the mouse E12.5 retina. The thick black bar depicts the mouse ECR9 region, the grey bars represent the regions of homology to the chicken, and the thin black bars represent motifs identified in the mouse ECR9 sequence.

**Additional File 7. Sequence alignments of ECR9 and ECR65 mouse, chicken and human homologous sequences.** Asterisks below nucleotides denote conservation. Labelled black arrows demarcate boundaries of Motifs or Regions that were deleted in Figure 4. ECR65 Region 3 and ECR65 Region 5 share a boundary. All deletions are directional as shown in Figure 4. Mutated bHLH sites are shown below full alignments, highlighted in blue.

**Additional File 8. Unscaled values from deletion, mutation, and overexpression experiments** (A) ECR65 activity from deletions and mutations corresponding to Figure 4. SP52 and NJ849 refer to two different orientations of ECR65::GFP. (B) ECR65 activity with empty pCAG vector, corresponding to Figure 5. (C) ECR9 activity from deletions and mutations, corresponding to Figure 4. NJ1140 and NJ1142 refer to two different orientations of ECR9. (D) ECR9 activity with the empty pCAG vector, corresponding to Figure 5. (E,F) Mutations of ECR65 and ECR9 with different mutant sequences, corresponding to Figure 4. Error bars represent 95% confidence interval. Each depicted point represents a biological replicate.

**Additional File 9 - Candidate bHLH factors are not sufficient to induce ectopic ThrbCRM1 activity in the mouse postnatal retina.** P0 mice retinae were electroporated with CAG::Bgal (magenta), ThrbCRM1::GFP (green) and the five candidate bHLH factors under the control of CAG. An empty CAG plasmid served as the negative control and CAG::OC1 as a positive control. Retinae were cultured for two days prior to harvest and staining with Bgal, GFP and DAPI.

**Additional File 10 - Individual bHLH factors are not sufficient to increase numbers of early retinal cell types.** E5 chick retinae were electroporated with CAG::bHLH constructs and CAG::Bgal as an electroporation control and cultured for two days before being processed for immunohistochemistry. Retinal sections were stained with DAPI (nuclei), Bgal (electroporated cells) and cell-type specific markers. Percentages were calculated using the number of cells marked by each factor out of the total number of Bgal(+) cells. Error bars represent 95% confidence intervals. Each point represents a biological replicate.

### Additional File 1

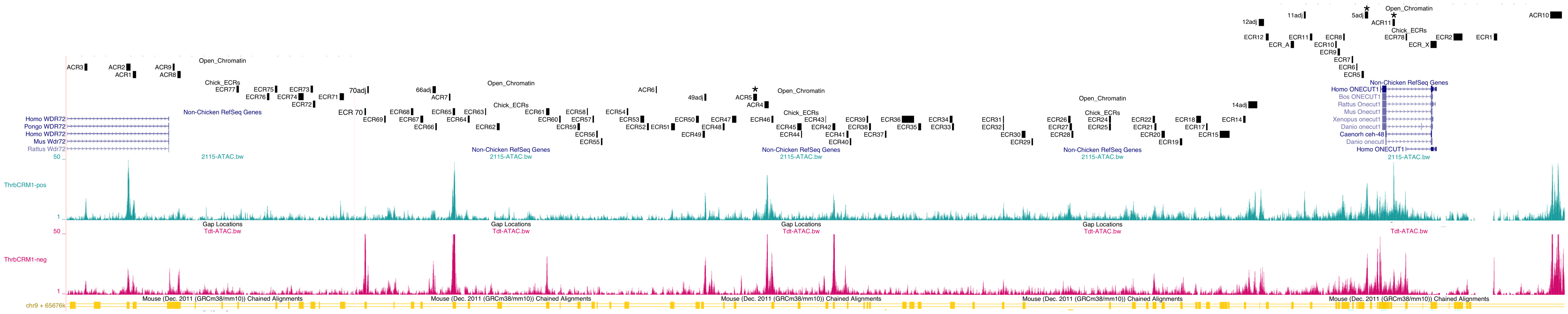

### Additional File 2

| Enhancer | Genomic Coordinates |
|---|---|
| **ECR1** | chr10: 8663469-8664114 |
| **ACR1** | chr10: 8385494-8386148 |
| **ECR2** | chr10: 8655229-8657051 |
| **ACR2** | chr10: 8384185-8385075 |
| **ECR_X** | chr10: 8650489-8651719 |
| **ACR3** | chr10: 8375625-8376231 |
| **ACR4** | chr10: 8514487-8515384 |
| **ECR5** | chr10: 8636448-8636927 |
| **ACR5** | chr10: 8512198-8513015 |
| **5adj** | chr10: 8637057-8637759 |
| **ECR6** | chr10: 8635344-8635535 |
| **ACR6** | chr10: 8492309-8492485 |
| **ECR7** | chr10: 8634455-8634704 |
| **ACR7** | chr10: 8450103-8450313 |
| **ECR8** | chr10: 8632688-8633007 |
| **ACR8** | chr10: 8394607-8395238 |
| **ECR9** | chr9:74846723-74847170 |
| **ACR9** | chr10: 8393675-8394030 |
| **ECR10** | chr10: 8631067-8631367 |
| **ACR10.A** | chr10:8675208-8675805 |
| **ACR10.B** | chr10:8675786-8676296 |
| **ACR10.C** | chr10:8676249-8676699 |
| **ACR10.D** | chr10:8676652-8677308 |
| **ECR11** | chr10: 8625901-8626415 |
| **ACR11** | chr10: 8642720-8643265 |
| **11adj** | chr10: 8624685-8625065 |
| **ECR_A** | chr10: 8622011-8622605 |
| **ECR12** | chr10: 8616893-8617477 |
| **12adj** | chr10: 8615429-8616540 |
| **ECR14** | chr10: 8612329-8612808 |
| **14adj** | chr10: 8613306-8615161 |
| **ECR15** | chr10: 8607507-8609556 |
| **ECR17** | chr10: 8604723-8605057 |
| **ECR18** | chr10: 8602677-8603712 |
| **ECR19** | chr10: 8599411-8599875 |
| **ECR20** | chr10: 8595596-8596293 |
| **ECR21** | chr10: 8594202-8594551 |
| **ECR22** | chr10: 8593808-8594297 |
| **ECR24** | chr10: 8584888-8585254 |
| **ECR25** | chr10: 8584888-8585254 |
| **ECR26** | chr10: 8576532-8577086 |
| **ECR27** | chr10: 8576893-8577385 |
| **ECR28** | chr10: 8577272-8577677 |
| **ECR29** | chr10: 8569146-8569350 |

| | |
|---|---|
| **ECR30** | chr10: 8567124-8567852 |
| **ECR31** | chr10: 8563261-8563496 |
| **ECR32** | chr10: 8563261-8563496 |
| **ECR33** | chr10: 8552799-8553199 |
| **ECR34** | chr10: 8552380-8552892 |
| **ECR35** | chr10: 8545908-8546480 |
| **ECR36** | chr10: 8542616-8545100 |
| **ECR37** | chr10: 8539165-8539457 |
| **ECR38** | chr10: 8535862-8536250 |
| **ECR39** | chr10: 8535401-8535717 |
| **ECR40** | chr10: 8531924-8532238 |
| **ECR41** | chr10: 8531318-8531625 |
| **ECR42** | chr10: 8528440-8528938 |
| **ECR43** | chr10: 8526952-8526952 |
| **ECR44** | chr10: 8521966-8522213 |
| **ECR45** | chr10: 8521266-8522058 |
| **ECR46** | chr10: 8515926-8516305 |
| **ECR47** | chr10: 8507915-8508786 |
| **ECR48** | chr10: 8505975-8506397 |
| **ECR49** | chr10: 8501851-8502327 |
| **49adj** | chr10: 8502221-8502639 |
| **ECR50** | chr10: 8500530-8501106 |
| **ECR51** | chr10: 8495478-8496218 |
| **ECR52** | chr10: 8490472-8490869 |
| **ECR53** | chr10: 8489091-8489919 |
| **ECR54** | chr10: 8486341-8486607 |
| **ECR55** | chr10: 8481081-8481361 |
| **ECR56** | chr10: 8480144-8480410 |
| **ECR57** | chr10: 8479342-8479641 |
| **ECR58** | chr10: 8478198-8478383 |
| **ECR59** | chr10: 8476275-8476779 |
| **ECR60** | chr10: 8472598-8472844 |
| **ECR61** | chr10: 8469909-8470514 |
| **ECR62** | chr10: 8459873-8460361 |
| **ECR63** | chr10: 8457431-8457602 |
| **ECR64** | chr10: 8453886-8454273 |
| **ECR65** | chr10: 8450796-8451312 |
| **ECR66** | chr10: 8447307-8447542 |
| **66adj** | chr10: 8446729-8447361 |
| **ECR67** | chr10: 8444225-8444787 |
| **ECR68** | chr10: 8442354-8442820 |
| **ECR69** | chr10: 8436831-8437103 |

| | |
|---|---|
| **ECR70** | chr10: 8432829-8433099 |
| **70adj** | chr10: 8433359-8433724 |
| **ECR71** | chr10: 8427766-8428649 |
| **ECR72** | chr10: 8422264-8422774 |
| **ECR73** | chr10: 8421785-8422274 |
| **ECR74** | chr10: 8419319-8420332 |
| **ECR75** | chr10: 8414515-8414968 |
| **ECR76** | chr10: 8412870-8413389 |
| **ECR77** | chr10: 8406698-8407162 |
| **ECR78** | chr10: 8645415-8645691 |

### Additional File 3

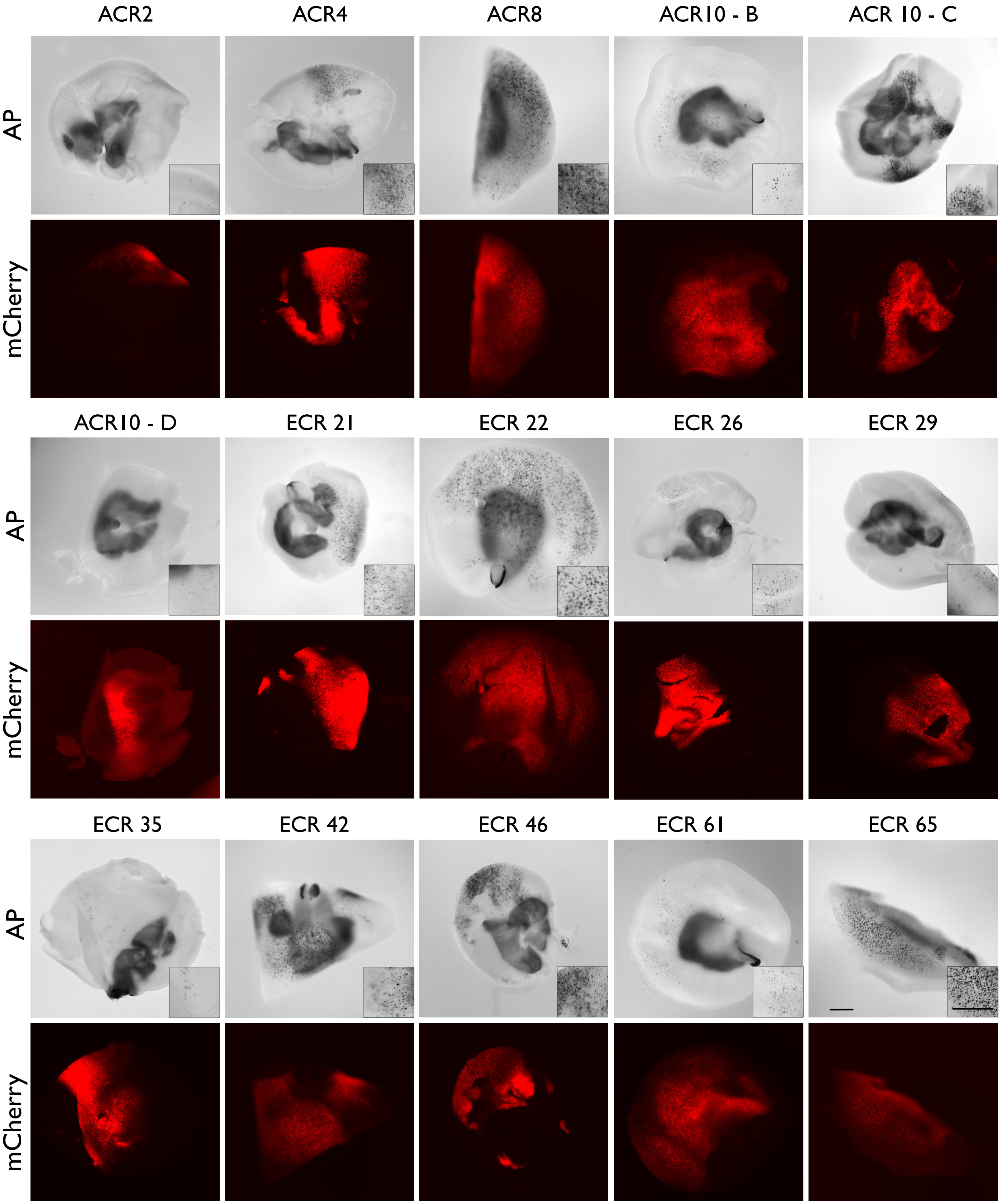

### Additional File 4

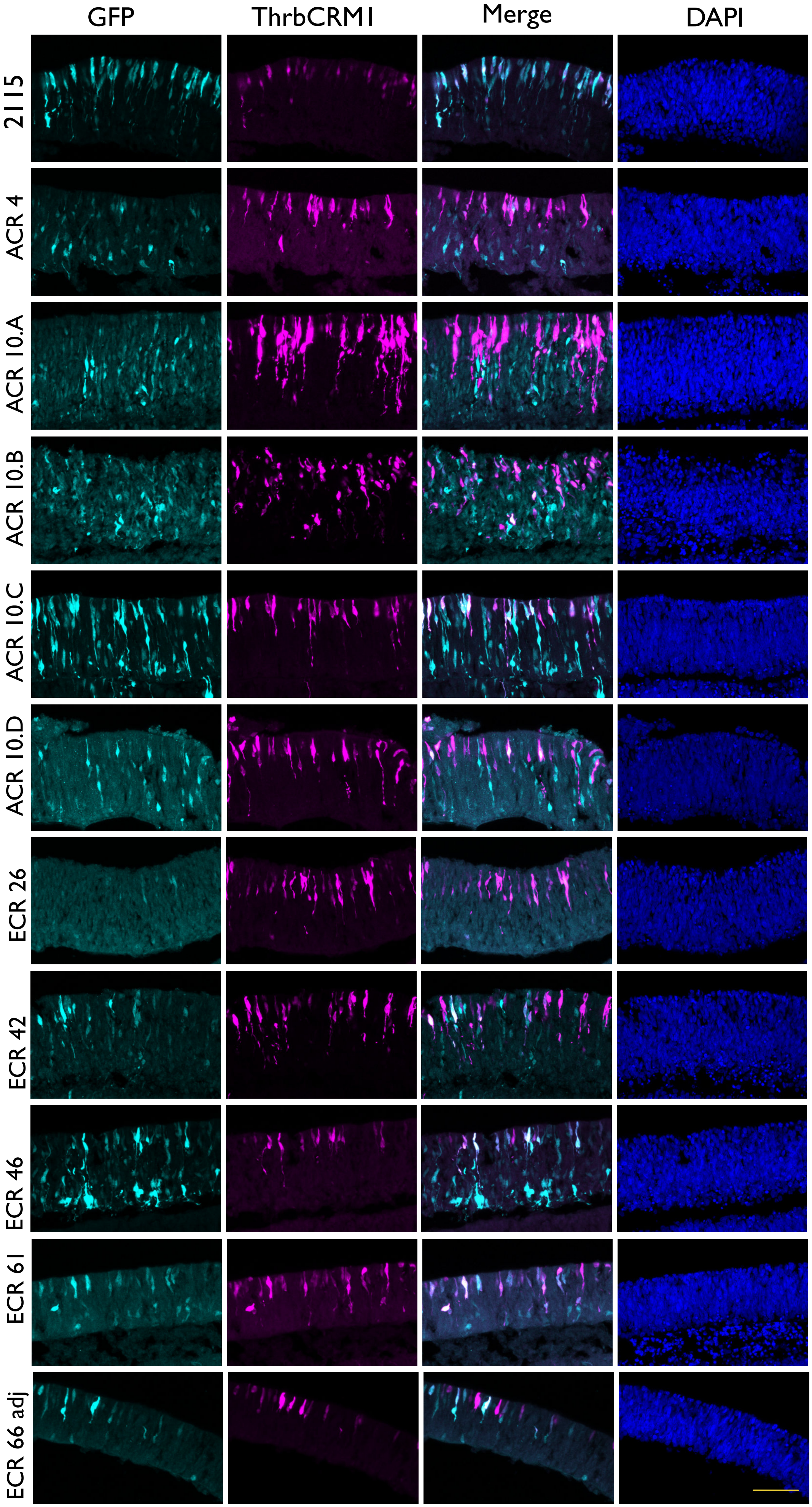

Additional File 5

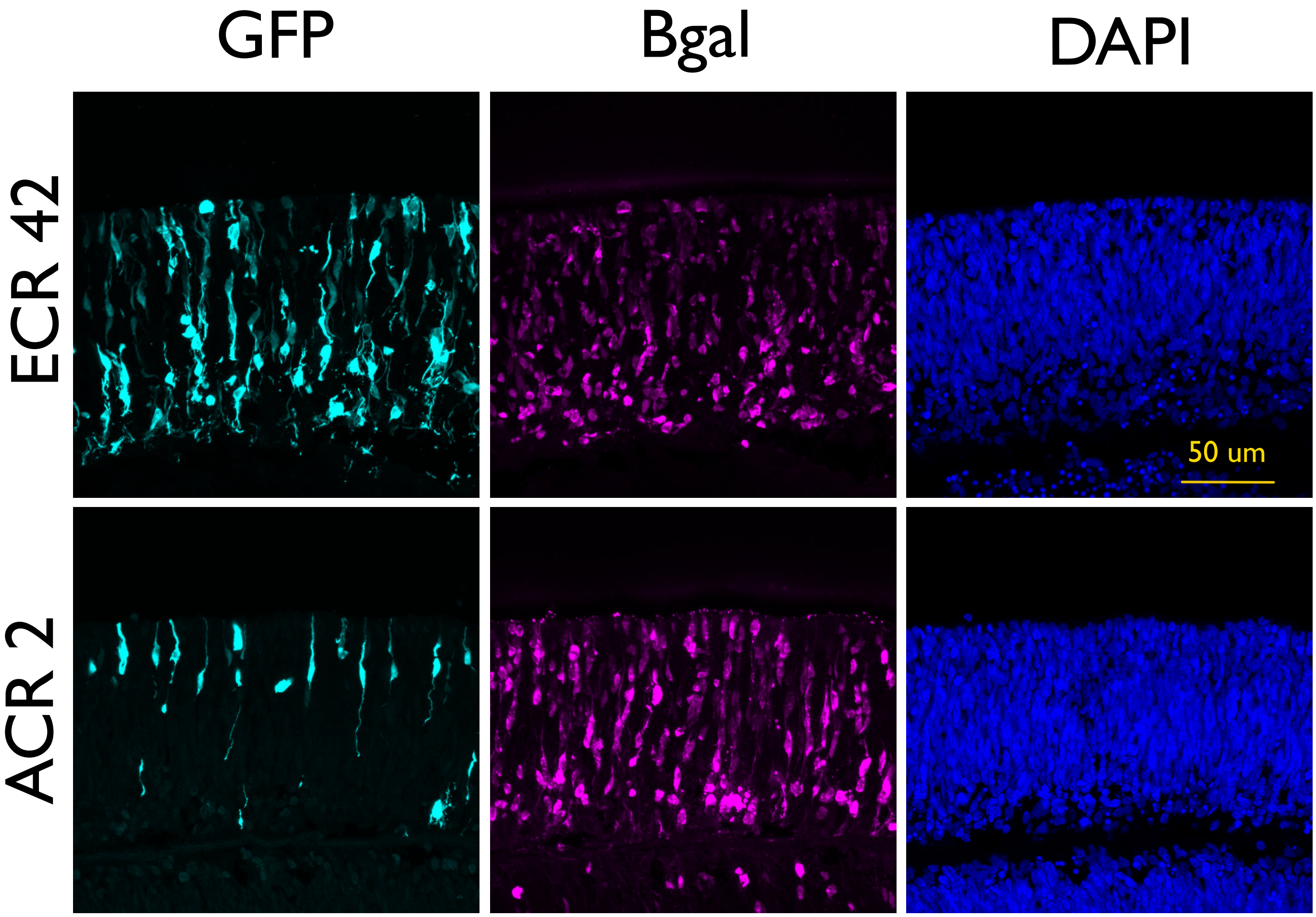

### Additional File 6

A

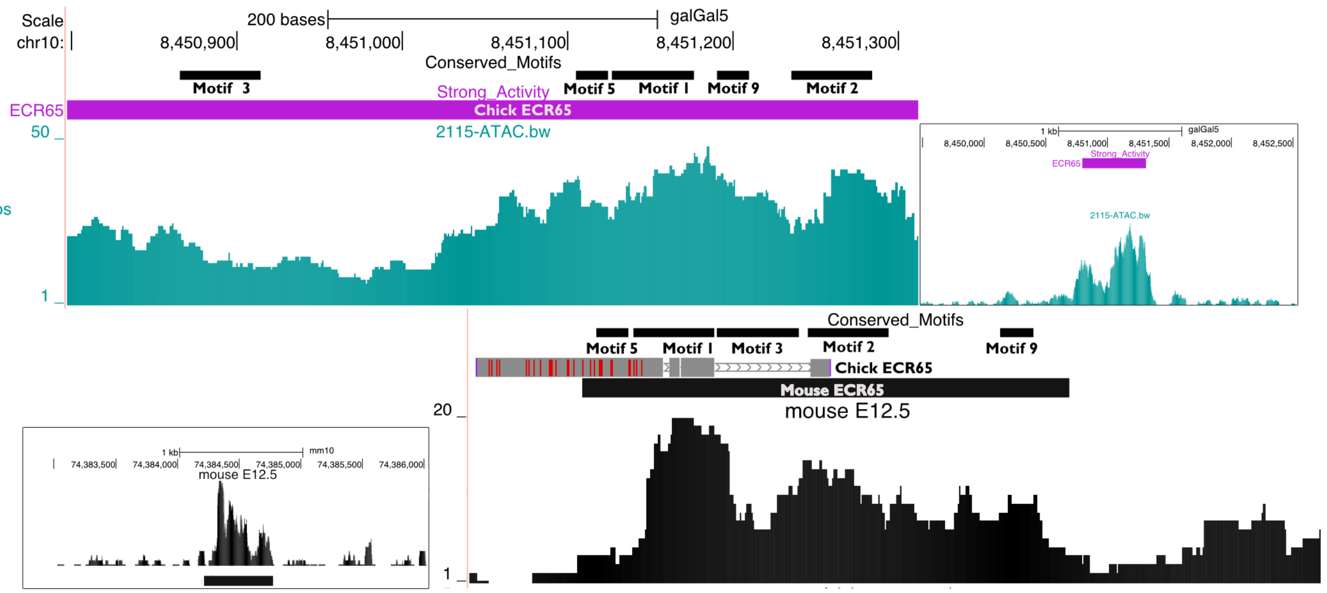


B

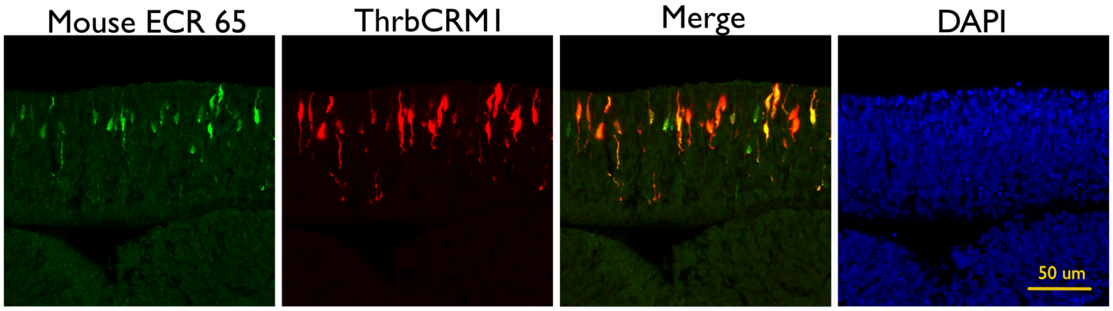


C

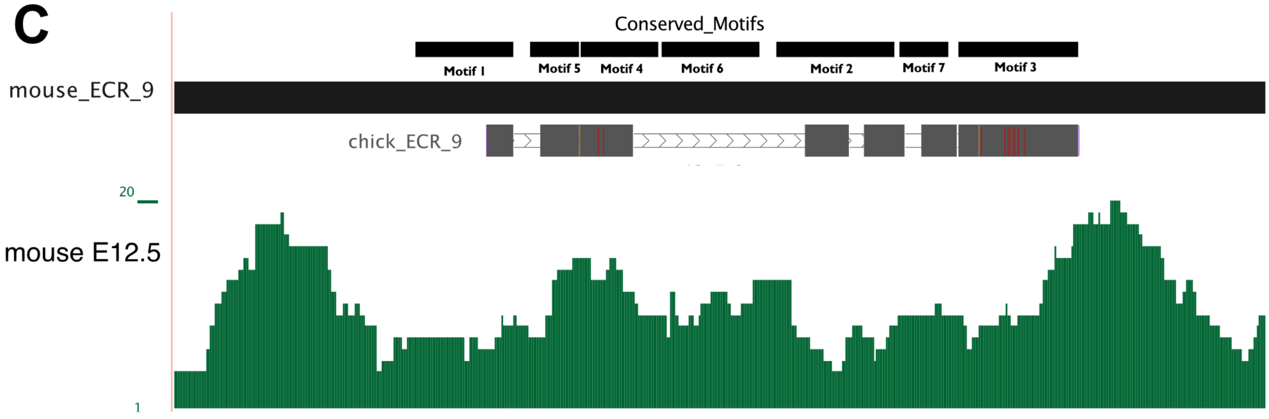

### Additional File 7

```
ECR9_chick ------------------------------------------------------------
ECR9_mouse TTGCACTTCCTGTGGATTAGTGAACAGATCTCTTTATAGGACTACATCTGTCTCTCTCTT
ECR9_human ------------------------------------------------------------

                                             Region F
ECR9_chick --------------------AGGGTAGGGCTTCCCAGGAAGGGTTAAGGCTTTTCAGGCC
ECR9_mouse CCACAATCTGTCCTACCCTAGGGGGAGGGCTGGCGGGAAAGGGTTAAAGGTTTTCTGGTC
ECR9_human ------------------------------------GAAAGGGTTAAAGGTTCTCTGGTC
                                           * ********* * ** ** ** *
                         Motif 1
ECR9_chick AAAATCCTCCAACAGAAGGCAGGGTCAGCCTCGGAGCTGAAATCTAAGTGCTAATGACTG
ECR9_mouse GAAATAGCCCAACAGA---AGGGGCCCGGCCCAGAGCTGAAATCTAAGT-CTAATGACCG
ECR9_human AAAATAGCCCAACAGA---AGGAGCTGGGCCCAGAGCTGAAATCTAAGTGCTAATGACCG
            ****   ********     * *   * * * *************** ******** *

ECR9_chick ATGCCCTTCACCCCTCCGTCGCCTCCAAAGGCGACTGCCCTGGATGAGC----CGGG---
ECR9_mouse CTGCCCTTCACCTCGCTGCTGTCCACAGGCCGGACCTGCGGCCCGGTGGGGGCCGGGACC
ECR9_human TTGCCCTTCACCCTACTGCTGTCCACTGGCCGGACCTGCGGCCGGGTGGGGCCCCGGACT
            ***********  * *  * *  *      ***   *      * *    * **
               Motif 2
ECR9_chick ------------GCCGGAGAGCATTTAACATTTGAACTTCTTCACGTCTGCGAGGCAGCA
ECR9_mouse AGGACAGGACACAATGTTCTTCGTTTAACATTTGAACTTC---TCCTATCCGAGGCAGCA
ECR9_human GGGAGAGGGCACAATATTCTGCATTTAACATTTGAACTTC---TCCTATCCGAGGCAGCA
                                * *****************    * * * *********
      Motif 7                                 Motif 3
ECR9_chick GGCCGTTGTGCGCGGGGT-CGCTGACCCCCAGCCCCGCTCCCAGCCCCTCAGCTGCCTC
ECR9_mouse GGCCGTTTTGTCCTGGGGTCGCTGAC----------CCTCGCTCCCCCATCAGCTGCCCT
ECR9_human GGCCGTTTTGTCCTGGGGTCGCTGAC----------CCTGGCTCCCCCATCAGCTGCCCT
            ******* **  * ***  *******           **  *    *** *********
                                    Region R
ECR9_chick CCGGCCCTAATTGAATGACAGATGGTCAGCGGCAGCGAGAGGGCAGGAGCCGGCCCCGCG
ECR9_mouse GAAACCTTAATTGAATGACAGATGGTCATTTCGGGGGCGCCAGAAGCTGCGCTGGCCAG
ECR9_human GAAACCTTAATTGAATGACAGATGGTCAT-------------------------------
               ** ********************

ECR9_chick CCCCCAGCACTGCTGCCCACGCGCCCCAGGGCCGCCTCGGCCGCCCACTAAGGGACGGGG
ECR9_mouse TGG-AGGGGCTCAGGCCGAGAAGGCCTTGCTTAGGCTCTGCCTGCT--------------
ECR9_human ------------------------------------------------------------
```

#### ECR9 bHLH Site 4

```
GGCCCTAATTGAATGACAGATGGTCAGC
AACCTTAATTGAATGACAGATGGTCATT
AACCTTAATTGAATGACAGATGGTCAT-
  ** *********************
```

#### ECR9 bHLH Site 3

```
CCGCTCCCCAGCCCCTCAGCTGCCTC
--CCTCGCTCCCCCATCAGCTGCCCT
--CCTGGCTCCCCCATCAGCTGCCCT
   **  *    *** *********
```

```
ECR65_chick TTTCCATGCATCAGCTGCTATTGTATGACCATACATAATTGGGTCTAAGTGCTTACATCA
ECR65_mouse ------------------------------------------------------------
ECR65_human ------------------------------------------------------------

ECR65_chick GATGCCTATTATGTGAAATTAATTTTTGGAACATATGCTGTTGTATTTATACCATTGTTG
ECR65_mouse ------------------------------------------------------------
ECR65_human ------------------------------------------------------------

ECR65_chick TTTCATGCTAGATTAATTCTTCAAGGGTTATAATCACATGACACTTATCATTTTAATTAT
ECR65_mouse ------------------------------------------------------------
ECR65_human ------------------------------------------------------------
                                     Region 1
ECR65_chick GCCTCTTTAGTTGCAACATATGACAGCTTTTGACTTCTCACCCTCTCCTTCTGCATGTGT
ECR65_mouse ---------------------------------------------------TTGTATTGTGT
ECR65_human ---------------------------------------------------TTGTATTGTGT
                                                            *     *****
                                     Region 2
ECR65_chick GACAGATAAAATTATTAATAAATGCTTGTTTATTATGTAGTTATGCTGTACATGTCCACA
ECR65_mouse GACCAATGAGATTATTAATAAATGCCTTTTCATTGTGTATCCAGGCTGTAGGTGCCCACG
ECR65_human GACCAATGAGATTATTAATAAATGCTTTTTTATTATGTAGTCATGCTGTAGGTGCCCACA
            ***  ** * *************** * ** *** ****   * ******  ** ****
                                                               Region 3/5
ECR65_chick GGGTGTAGGTTGGCCTTGTGTATTTTACCCAGAGTACAGTGTCAGCAGAGCCTTTGTGGT
ECR65_mouse GGGTATGGGGCAGCCTCCTGTATTTTATTCTGGGTGCAGTGTCAGCAGGACCTTGTGGAT
ECR65_human GGGTATGGGCTGGCCTCATGTATTTTATTCTGGGTGCAGTGTCAGCAGGGCCTTGCGGAT
            **** * **   ****  *********  * * ** ************  ****  * *
                                                               Region 4
ECR65_chick TTGCTAATGAGCTAAATCCCACTAACACCACACACACTCTAAACTGCCAAAAGCCCTTTA
ECR65_mouse TTGCTAATGAGCTAAATCCGGTTAGTGCCACACGCCCTGGATGGC--CAGAGATCTTTTA
ECR65_human TTGCTAATGAGCTAAATCCGGTTAGTGCCACATGCCCTGGACTGC--CAGAGATCTTTTA
            *******************   **   *****  * **  *       ** *   * ****

ECR65_chick CACATTCAAACACATACAAAGACTCAGCTGGGTCCTTTGTATAAAAGATTAACTCTTTAT
ECR65_mouse AATACATCATTGCTCACAGAGACTCAGCTGGACTCAGGCAGAGAGAGATTAACTCCTCAT
ECR65_human AATACATCATTGCTCTCAGAGACTCAGCTGGGTTCAGTATGAGAGAGAGATTAACTTCTCAT
             * *    *   *    ** ************    *        * *********  * **

ECR65_chick TCCAATTAAAGATGAGCTGGCCTGGCTTGGTCCAA-------------------------
ECR65_mouse TCCAATTAGCAAAGAGTCCGACTGTTGCATGT-GGCTTGCTTCTGTAGCCCCAAACCCGG
ECR65_human TCCAATTAACAGAGAGTCATACTGCTGGAGAGAGGGTTGCTTATGTAGCCCCAAACCTAG
            ********      ***      ***
```

#### ECR65 bHLH site

```
TACAAAGACTCAGCTGGGTCCTT
CACAGAGACTCAGCTGGACTCAG
CTCAGAGACTCAGCTGGGTTCAG
 ** ************    *
```

### Additional File 8

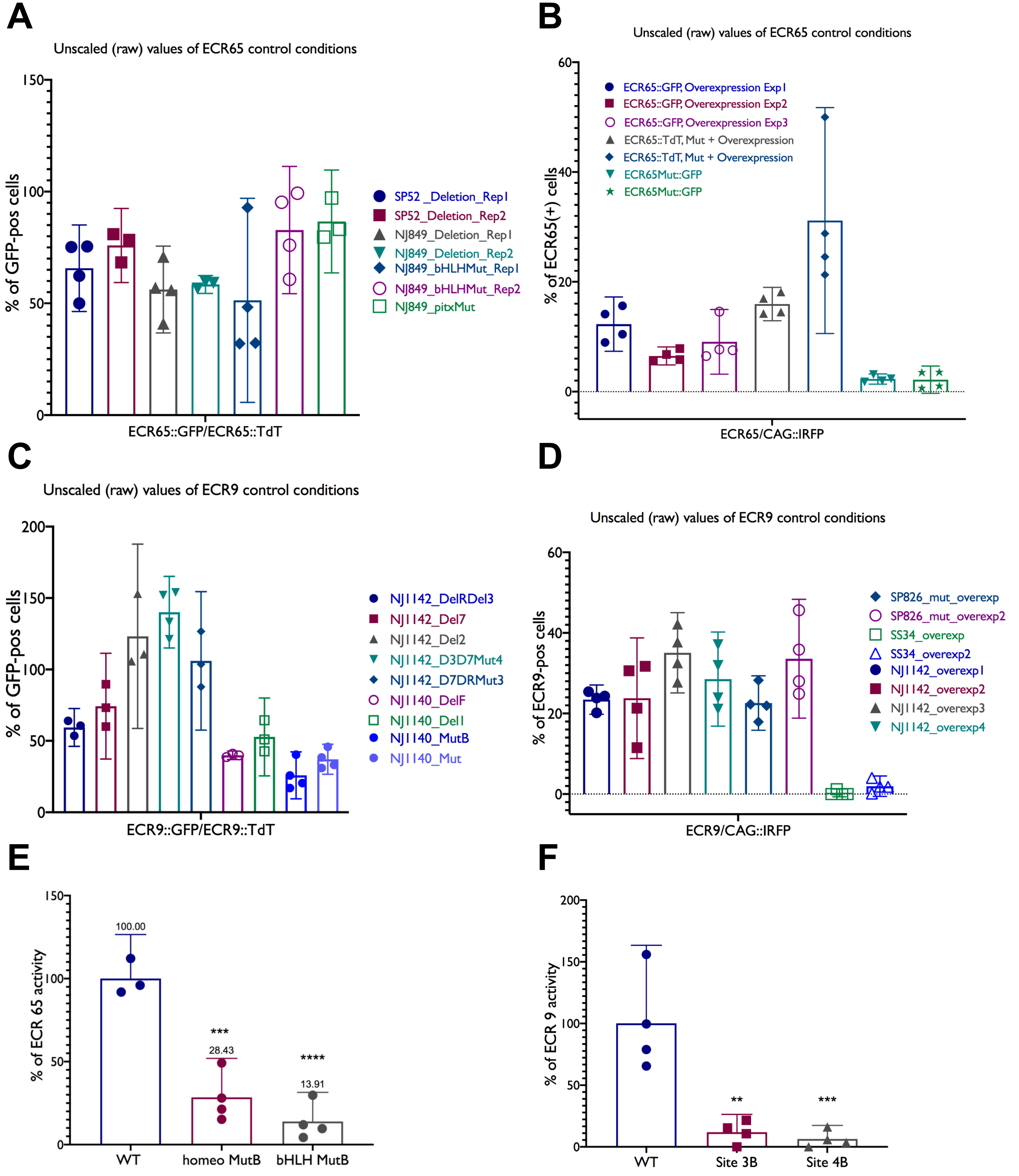

### Additional File 9

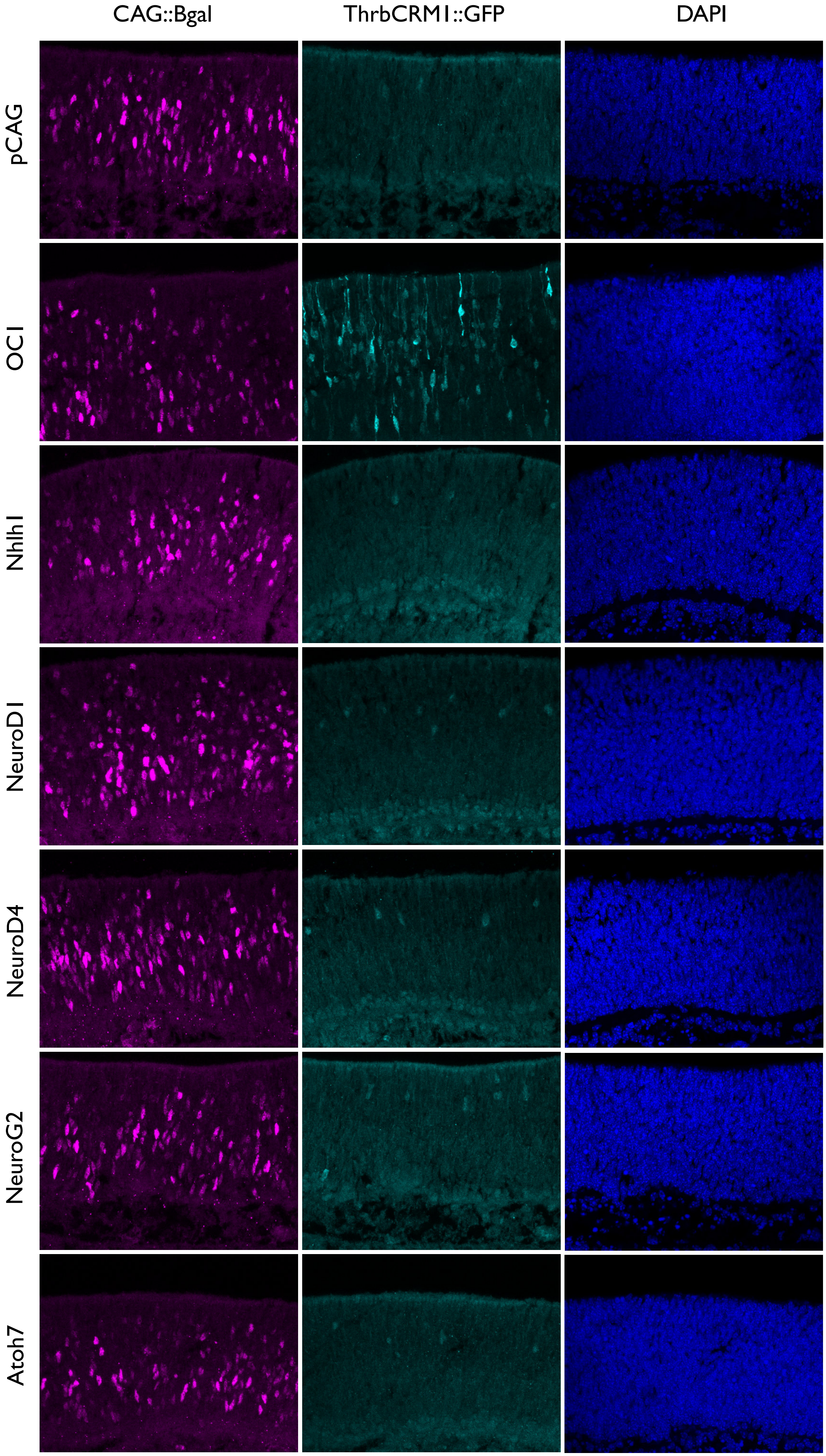

### Additional File 10

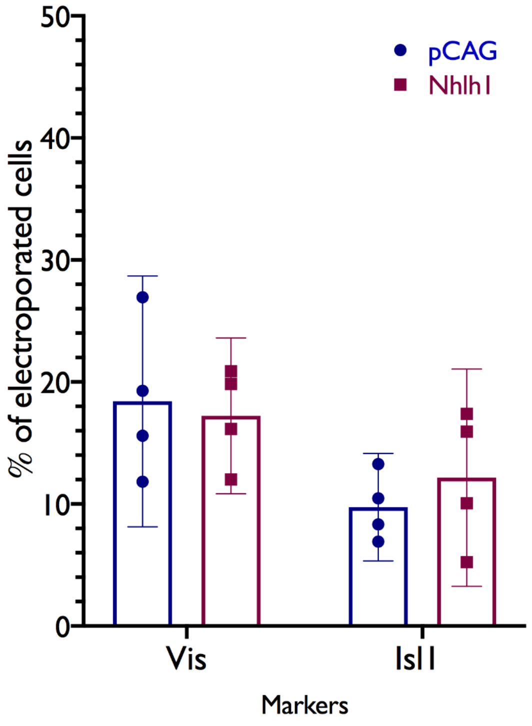


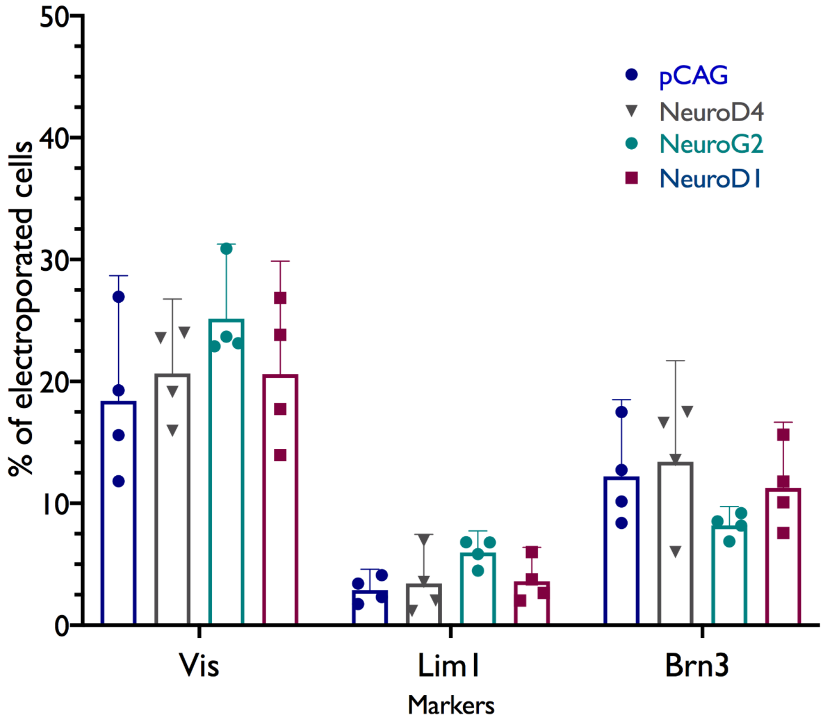
